# Alpha-synuclein regulates the repair of genomic DNA double-strand breaks in a DNA-PK_cs_-dependent manner

**DOI:** 10.1101/2024.02.29.582819

**Authors:** Elizabeth P. Rose, Valerie R. Osterberg, Jovin S. Banga, Vera Gorbunova, Vivek K. Unni

## Abstract

α-synuclein (αSyn) is a presynaptic and nuclear protein that aggregates in important neurodegenerative diseases such as Parkinson’s Disease (PD), Parkinson’s Disease Dementia (PDD) and Lewy Body Dementia (LBD). Our past work suggests that nuclear αSyn may regulate forms of DNA double-strand break (DSB) repair in HAP1 cells after DNA damage induction with the chemotherapeutic agent bleomycin^1^. Here, we report that genetic deletion of αSyn specifically impairs the non-homologous end-joining (NHEJ) pathway of DSB repair using an extrachromosomal plasmid-based repair assay in HAP1 cells. Importantly, induction of a single DSB at a precise genomic location using a CRISPR/Cas9 lentiviral approach also showed the importance of αSyn in regulating NHEJ in HAP1 cells and primary mouse cortical neuron cultures. This modulation of DSB repair is dependent on the activity of the DNA damage response signaling kinase DNA-PK_cs_, since the effect of αSyn loss-of-function is reversed by DNA-PK_cs_ inhibition. Using *in vivo* multiphoton imaging in mouse cortex after induction of αSyn pathology, we find an increase in longitudinal cell survival of inclusion-bearing neurons after Polo-like kinase (PLK) inhibition, which is associated with an increase in the amount of aggregated αSyn within inclusions. Together, these findings suggest that αSyn plays an important physiologic role in regulating DSB repair in both a transformed cell line and in primary cortical neurons. Loss of this nuclear function may contribute to the neuronal genomic instability detected in PD, PDD and DLB and points to DNA-PK_cs_ and PLK as potential therapeutic targets.

## Introduction

Synucleinopathies, such as Parkinson’s Disease (PD), Parkinson’s Disease Dementia (PDD), and Lewy Body Dementia (LBD), are characterized by the presence of aggregated α-synuclein (αSyn) pathology, known as Lewy pathology, which is found in surviving neurons in brain regions vulnerable to cell death. Although these three synucleinopathies, in addition to several other neurodegenerative diseases, are characterized by this abnormal αSyn aggregation within cell bodies and neurites, known as Lewy bodies and neurites, respectively, it is still not clear how Lewy pathology relates to neuronal cell death. Evidence exists for both gain-of-function and loss-of-function mechanisms for how αSyn aggregation may play a role. Possible gain-of-function mechanisms include toxic αSyn aggregates taking on distinctly new properties and disrupting presynaptic^2,3^ & mitochondrial function^4^, affecting protein degradation by the ubiquitin-proteasome system^5,6^ or autophagy^7–9^.

αSyn is normally localized to the presynaptic terminal and nucleus of neurons, which is how it was originally named^10^. It is a 140 amino acid long protein that binds to the outer leaflet of synaptic vesicles and contributes to their trafficking^11^. Extensive work has characterized its normal physiological function in modulating vesicle sorting and clustering at presynaptic terminals^12^ during exo- and endocytosis^2^. Presynaptic loss-of-function hypotheses for the role of αSyn in synucleinopathies suggest dysregulation of these neurotransmitter vesicle processing steps in neurons with Lewy inclusions. Conflicting evidence exists for whether αSyn knockdown is protective^13^ or harmful^14^ in PD models. αSyn is also found in the nucleus of cells^15–17^, where its role is less clear. Selective localization of αSyn to the nucleus using a nuclear localization sequence has been associated with motor deficits independent of its aggregation in mice^18^. Other evidence points to the importance of αSyn in regulating transcription, modifying histone biology, and directly binding DNA^19–21^. It has also be recently suggested to influence mRNA stability in P-bodies^22^. Our previous work highlights a normal physiologic role in regulating forms of DNA repair, including double-strand break (DSB) repair^1^, and was motivated, in part, by human pathological studies in patients with Ataxia-Telangiectasia (AT) showing that they can develop Lewy pathology^23^. AT is a result of defects in DSB repair caused by mutations in the Ataxia-Telangiectasia Mutated (ATM) gene, known to be a critical kinase involved in DNA damage response signaling after DSBs have been detected within a cell, where it then helps to coordinate repair. An important aspect of this signaling is the phosphorylation of the histone H2AX, creating a mark known as γH2AX that helps recruit downstream DSB repair components to the site of the break. Interestingly, work in a mouse ATM knock-out (KO) model also suggests that this DSB repair dysfunction can cause progressive dopamine neuron loss in the substantia nigra and αSyn aggregation similar to what is seen in people with PD^24^.

Our previous work demonstrated that the induction of DNA damage with the chemotherapeutic agent bleomycin in the leukemia-derived immortalized HAP1 cell line clearly increased DSB and γH2AX levels, and that this increase was even greater in the αSyn KO condition. Bleomycin is used extensively to chemically induce DSBs in multiple systems and this has been shown to be associated with increased γH2AX levels, however, a caveat of this approach is that bleomycin is also known to increase other forms of DNA damage besides DSBs^25,26^. Importantly, although γH2AX is widely used as a marker of DSB repair signaling, it can also be increased by other forms of DNA damage like clustered oxidative lesions or high levels of single-strand breaks^27^. In order to directly test the role of αSyn in the repair of DSBs specifically, we set out here to use CRISPR/Cas9-based methods to selectively induce a single DSB at a prespecified site in the human genome in HAP1 cells to test the normal role of αSyn in the DSB repair process. In addition, we also set out to use this CRISPR/Cas9 strategy to induce a similar, prespecified DSB in the genome of primary cortical neurons from mice to test whether αSyn is important for modulating DSB repair in this cell type as well.

Our recent work studying the ability of αSyn to bind double-stranded DNA using an *in vitro* gel-shift assay suggested that it can directly bind in the major groove. Interestingly, phosphorylation of αSyn at serine-129 (S129) seemed to alter the properties of this binding and switched DNA binding to a different mode^28^. S129 phosphorylation (pSyn) is the most common post-translational modification αSyn undergoes and is highly enriched in Lewy pathology^29^. It is unclear, however, whether pSyn promotes or protects against aggregation^30^. While S129 phosphorylation has been shown to reduce αSyn fibrillogenesis *in vitro*^31^, other *in vitro* studies suggest that it can also promote fibrillar aggregation depending on the exact conditions used^32–34^. *In vivo*, studies utilizing a phospho-deficiency approach where S129 is mutated to alanine increased aggregation in drosophila^35^, but this same S129A mutant had no effect^36^ or reduced^37^ aggregation compared to the phospho-mimic S129D mutation in rat brain. Another strategy to study the effects of S129 phosphorylation is to modulate the kinases and phosphatases that act at this residue. Several kinase families have been shown to produce pSyn *in vitro*^38–42^, but several studies, including our own work, suggest that the Polo-Like Kinase (PLK) family members 1, 2, and 3^43–49^ may be the most important *in vivo*^50^. Our previous work demonstrated that genetic knockout of PLK2 led to increased αSyn in nuclear DSB repair foci and to improved survival of cortical neurons bearing aggregated αSyn inclusions in mouse cortex *in vivo*^50^. Given the attractiveness of kinases, specifically the PLK family and PI3KK family, important for DSB repair signaling (e.g. ATM, ATR, DNA-PK_cs_), as drug targets, we also set out here to test their potential relevance to nuclear αSyn biology in modulating DSB repair, genomic stability and neurodegeneration.

## Results

### αSyn KO in HAP1 cells impairs non-homologous end-joining

Our previous research in HAP1 cells suggested a role for αSyn in DSB repair using γH2AX levels as primary readout, however, there are two main mechanisms for repairing DSBs: non-homologous end-joining (NHEJ) and homologous recombination (HR). Perturbations in either can alter γH2AX levels. To directly test which pathway αSyn may be involved in, we used a previously established plasmid-based reporter system. This system involves two different plasmids, both of which can be linearized and transfected into cells. One is sensitive to NHEJ repair and will only produce fluorescent GFP expression when the plasmid is repaired by NHEJ and re-circularized in the process. An analogous plasmid assays HR and only produces functional GFP when HR repairs and re-circularizes this plasmid^51^. After transfection of these linearized plasmids into HAP1 cells, we measured GFP expression and RFP expression (as a transfection control) using flow cytometry at 72 hours post-transfection (Fig. 1A). We only found a significant difference between WT and αSyn KO cells when using a plasmid reporting NHEJ repair (Fig. 1B). This suggests that the loss of ⍺Syn impairs NHEJ repair efficiency. We did not find a significant difference between WT and αSyn KO cell lines with the HR reporter (Fig. 1B), although overall repair efficiency was lower with this plasmid, possibly making the assay less sensitive to HR repair differences.

**Figure 1.**
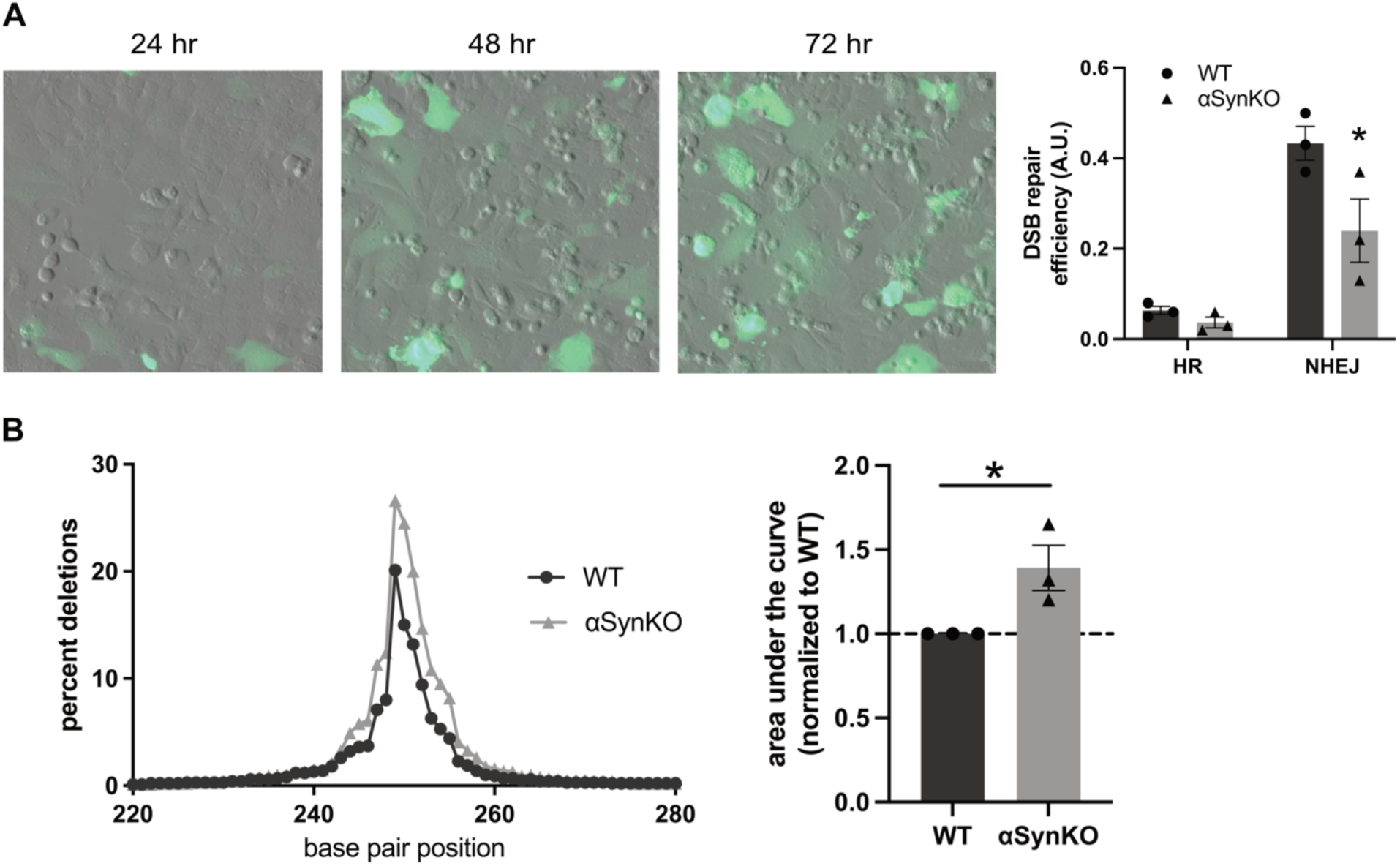
αSyn KO in HAP1 cells reduces NHEJ efficiency using a plasmid reporter system. A) NHEJ reporting green fluorescing HAP1 cells after repair event at 24, 48, and 72 hours post transfection (Scale bar = 50 μm). RIGHT: Quantification from 72 hour timepoint. HR reporter in WT cells (0.063 ±0.009) vs HR reporter in KO cells (0.037 ±0.012)(p=0.2697), NHEJ reporter in WT cells (0.433 ±0.038) vs KO cells (0.240 ±0.070)(p=0.0276). N=∼500,000 cells counted per replicate, 3 biological replicates. Two-tailed student’s t-test. C) Next Generation Sequencing of repair junction from WT and KO HAP1 cells transfected with NHEJ reporter. Percent of deletions are increased in KO cells compared to WT cells. RIGHT: Quantification. Area under the curve of KO cells (1.392 ±0.134) significantly increased compared to WT cells (1.0 ±0.0)(p=0.0435) Two-tailed student’s t-test.

One characteristic of NHEJ is that it is more error-prone compared to HR, since the repair that occurs is not templated using a faithful copy of the sequence from the sister chromatid, as it is in HR. This often leads to the introduction of insertions and deletions (indels) at the repaired junction. In addition, NHEJ can be broken down to different subtypes, with classical NHEJ (c-NHEJ) being the most well studied and dependent on specific components like DNA-PK_cs_, XRCC4/XLF and ligase 4^52^. c-NHEJ is thought to be the least error-prone of the NHEJ pathways, with a decreased likelihood of introducing indels during repair and smaller sized indels when they do occur. In addition to c-NHEJ, other alternative NHEJ (alt-NHEJ) pathways dependent on Pol *θ*, XRCC1, ligase 3, and single-strand annealing (SSA) pathways have been described. When alt-NHEJ is recruited to repair DSBs, higher levels and larger indel sizes are reported in the repaired products^53,54^. With this in mind, we next wanted to investigate how the loss-of-function of αSyn affects the size and frequency of indels at the DSB repair junction. In order to do this, we transfected a ∼500 bp linear double-stranded DNA into WT or αSyn KO HAP1 cells. DNA that had undergone DSB repair to produce a circular topology was purified from cells, then exposed to exonuclease treatment to remove remaining linear DNA from the sample, purified, linearized again (using a cut site opposite the repair junction) and analyzed by next-generation sequencing. This approach allowed us to detect the frequency of repair events that incorporated an indel during the repair process and to determine the exact size and position of these indels. Using this strategy, we found an increase in the frequency of sequenced junctions with deletions incorporated at the repair site in αSyn KO cells compared to WT cells (Fig. 1B). There was no apparent change in the size spectrum of deletions, however, and they were also relatively small (most <10 bp in length). This suggests that αSyn may be influencing specific sub-forms of c-NHEJ, since alt-NHEJ and SSA are usually associated with larger deletions (>20 bp in length)^55^. The repair of extrachromosomal DNA transfected into cells is likely, however, to involve different processes than occur in the context of genomic DSBs, where other important factors like DNA methylation, histone modifications, and higher order chromatin organization are important.

### αSyn KO modulates repair fidelity after CRISPR/Cas9-induced DSB formation in HAP1 cells

To assess whether αSyn is involved in NHEJ within this genomic DSB context, we turned to a CRISPR/Cas9 approach using lentiviral expression. Cells were transduced with lentivirus to introduce a single DSB in the *DNMT3B* locus. As a control for transduction efficiency, a similarly designed lentivirus expressing EGFP was used. WT and αSyn KO HAP1 cells were transduced by this lentivirus similarly when assayed using immunocytochemistry (Fig. 2A). In order to confirm this using a second approach, we also used flow cytometry to quantify transduction efficiency and no significant differences between WT and αSyn KO HAP1 cells were observed (Fig. 2B, see Methods).

**Figure 2.**
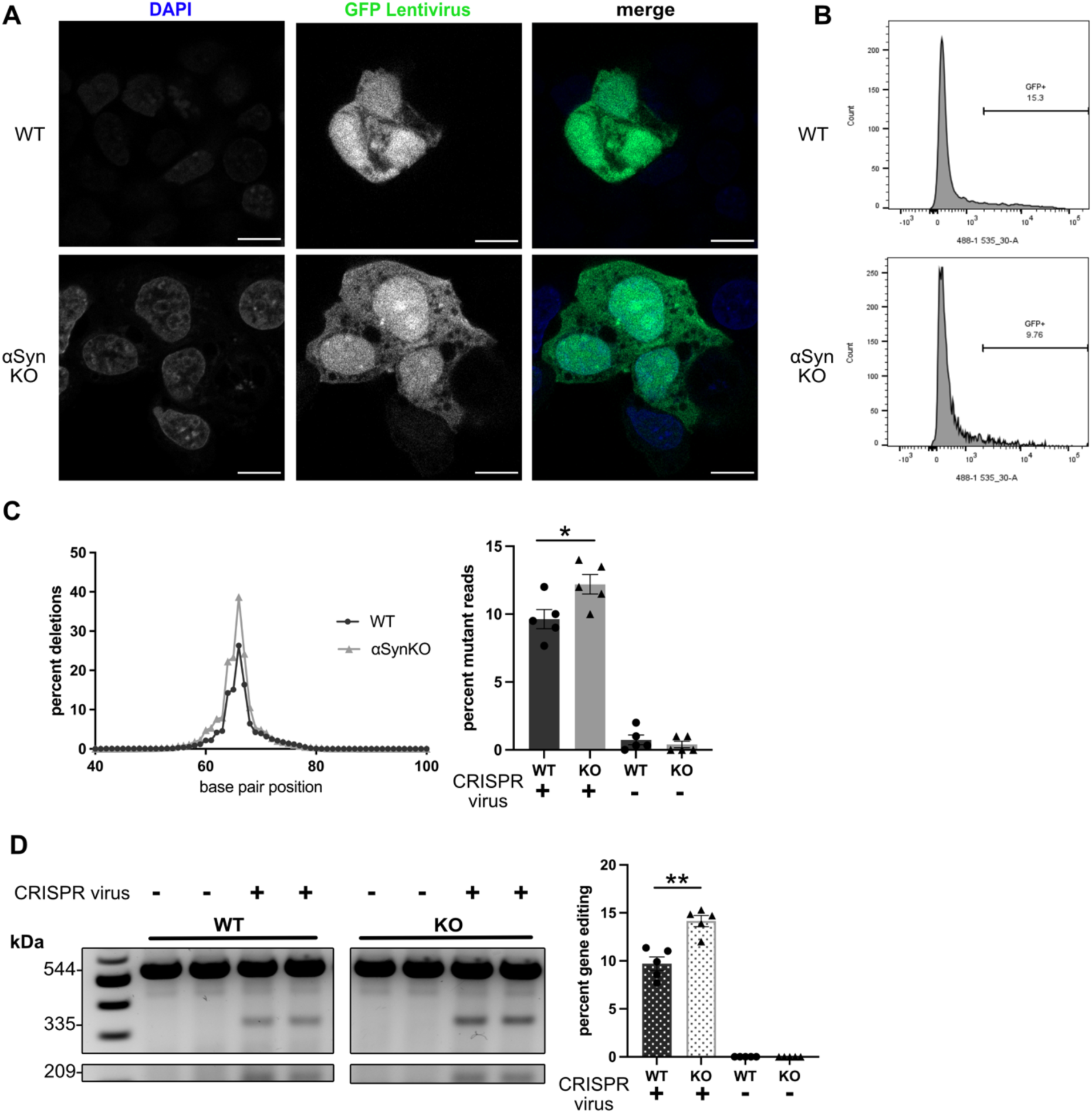
αSyn KO in HAP1 cells reduces genomic DSB NHEJ efficiency using a CRISPR/Cas9 system. A) GFP tagged DSB inducing CRISPR/Cas9 lentiviral treated WT and αSyn KO HAP1 cells. Scale bar 10μm. B) Representative histograms of GFP tagged lentiviral transduction efficiency in WT and αSyn KO HAP1 cells using flow cytometry. No significant difference observed (see methods). C) NGS of 288bp repair junction with percent of reads containing insertions (green) or deletions (blue) from WT and αSyn KO HAP1 cells transduced with CRISPR/Cas9 DSB inducing lentivirus. Mutant reads plotted against base pair position with cut site at base pair 67. RIGHT: Quantification. Increased mutant reads in αSyn KO cells (12.20 ±0.718) compared to WT cells (9.633 ±0.708)(p=0.0344). WT Nontargeting Virus (0.733 ±0.356), αSyn KO Nontargeting Virus (0.400 ±0.245). N=5 biological replicates (2-3 technical replicates per biological replicate). Two-tailed student’s t-test. D) T7 Endonuclease I enzymatic assay gel image of full length unedited amplicon 544bp, edited fragmented products 335bp and 209bp. RIGHT: Quantification. % Gene Editing = 100 x (1 – (1-fraction cleaved)^1/2^). Significant increase in gene editing in αSyn KO HAP1 cells (14.15 ±0.583) compared to WT cells (9.707 ±0.702)(p=0.0012). WT Nontargeting virus (0.0 ±0.0), αSyn KO Nontargeting Virus (0.0 ±0.0). N=5 biological replicates (2-3 technical replicates per biological replicate). Two-tailed student’s t-test.

To assess the effect of αSyn on the repair of single DSBs induced at the *DNMT3B* locus, we performed PCR to amplify a 288 bp product across the repair junction and analyzed these for indels using next-generation sequencing. Interestingly, similar to our data using transfected DNA (Fig. 1B), when we analyzed DSB repair junctions induced in genomic DNA we also found a significant increase in the frequency of repair junctions containing deletions in αSyn KO cells compared to WT cells (Fig. 2C), with little change in the spectrum of deletions induced and relatively small lengths (most <10 bp). This again suggests that the loss of ⍺Syn compromises DSB repair fidelity and may be skewing DSB repair between sub-forms of c-NHEJ that are more likely to induce small deletions at the junction. This finding was confirmed by a secondary method for measuring repair fidelity using a T7 Endonuclease 1 (T7EI) assay. In this case, we amplified a longer PCR product flanking the repair junction (544 bp) and digested the products with T7EI, which recognizes and cleaves duplex DNA that contain imperfect base pairing. These cleaved fragments of DNA are measured to determine the ratio of edited to unedited PCR products in the sample. This provides a way to screen for indels that were introduced during the repair of the CRISPR-mediated DSB and these results were similar to our sequencing data. We found a significant increase in gene editing (T7EI cleavable product) using this approach in αSyn KO cells compared to WT cells (Fig. 2D). Together, these two assays for indel frequency show that αSyn loss-of-function leads to an increase in small indel frequency during the repair of DSBs induced at an individual site in the human genome.

### αSyn KO affects repair fidelity after CRISPR/Cas9-induced DSB formation in mouse primary cortical neurons

We next wanted to investigate whether αSyn also affects DSB repair fidelity in neurons, where NHEJ is thought to be the primary mechanism for DSB repair. We cultured WT and αSyn KO E18 cortical mouse neurons *in vitro* and used an analogous *Dnmt3b* targeting CRISPR/Cas9 approach using lentiviral expression. We first measured transduction efficiency using lentivirus expressing EGFP in WT and αSyn KO neurons using immunocytochemistry and detected no significant differences (Fig. 3A, see Methods). We then measured the indel frequency after the repair of a single DSB introduced into the genome of WT and αSyn KO mouse neurons using a PCR amplification and sequencing assay similar to the one we used in HAP1 cells. We again found a significant increase in the fraction of repair junctions containing deletions in αSyn KO neurons compared to WT neurons (Fig. 3B), very similar to what we found in HAP1 cells (Fig. 2C). Interestingly, the deletions in neurons were smaller around the cut site (most <5 bp in length) than what we detected in HAP1 cells (Fig. 2C). We next tested whether loss of αSyn leads to increased indel frequency using the T7EI assay in cortical neurons and again detected an increase in gene editing in αSyn KO neurons compared to WT (Fig. 3C).

**Figure 3.**
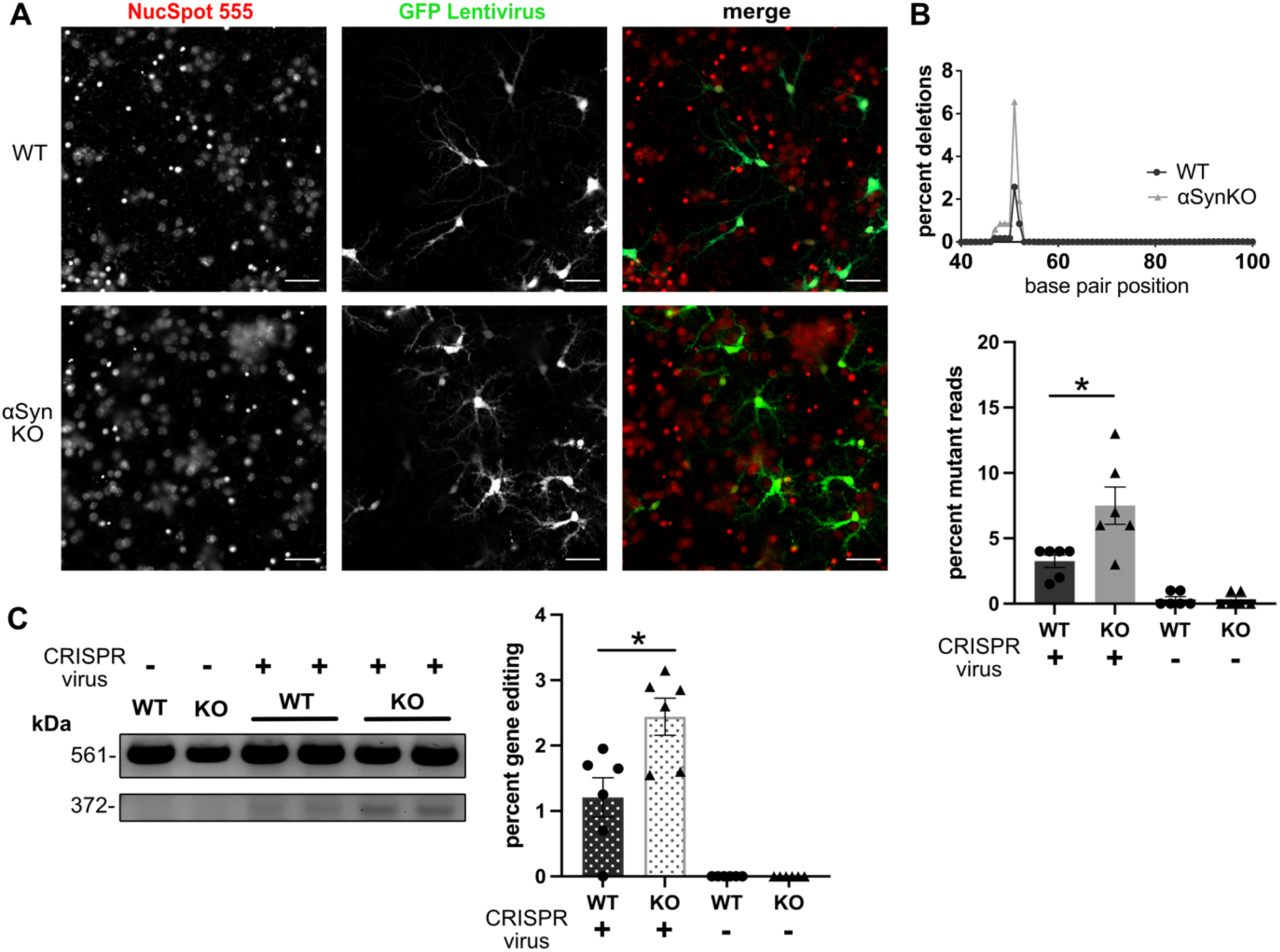
αSyn KO in mouse cortical neurons reduces genomic DSB NHEJ efficiency using a CRISPR/Cas9 system. A) GFP tagged DSB inducing CRISPR/Cas9 lentiviral treated WT and αSyn KO mouse E18 cortical neurons. Scale bar 20μm. B) NGS of 272bp repair junction with percent of reads containing insertions (green) or deletions (blue) from WT and αSyn KO mouse neurons transduced with CRISPR/Cas9 DSB inducing lentivirus. Mutant reads plotted against base pair position with cut site at base pair 52. RIGHT: Quantification. Increased mutant reads in αSyn KO neurons (7.500 ±1.568) compared to WT cells (3.250 ±0.524)(p=0.0183). WT Nontargeting Virus (0.333 ±0.231), αSyn KO Nontargeting Virus (0.333 ±0.231). N=6 biological replicates (1-2 technical replicates per biological replicate). Two-tailed student’s t-test. C) T7 Endonuclease I enzymatic assay gel image of full length unedited amplicon 561bp, edited fragmented product 372bp. RIGHT: Quantification. % Gene Editing = 100 x (1 – (1-fraction cleaved)^1/2^). Significant increase in gene editing in αSyn KO neurons (2.442 ±0.283) compared to WT cells (1.208 ±0.300)(p=0.0136). WT Nontargeting virus (0.0 ±0.0), ⍺Syn KO Nontargeting Virus (0.0 ±0.0). N=6 biological replicates (1-2 technical replicates per biological replicate). Two-tailed student’s t-test.

### αSyn modulates repair fidelity via DNA-PK_cs_-dependent mechanism

To test how αSyn interacts with polymerases, nucleases and kinases known to be important in various DSB repair pathways, we next set out to test how a panel of small molecule inhibitors interacted with DSB repair in αSyn WT and KO HAP1 cells using our CRISPR-mediated DSB induction approach. To test the roles of Polymerase *θ* and Mre11, we used pharmacological inhibitors of Polymerase *θ* (ART558) and Mre11 (mirin). We did not observe any consistent differences in the indel frequency after DSB repair in WT and αSyn KO HAP1 cells treated with either a Pol*θ* inhibitor (Pol*θ*i) or Mre11i, compared to a vehicle control using both our sequencing and T7EI assays (Fig 4A-B).

**Figure 4.**
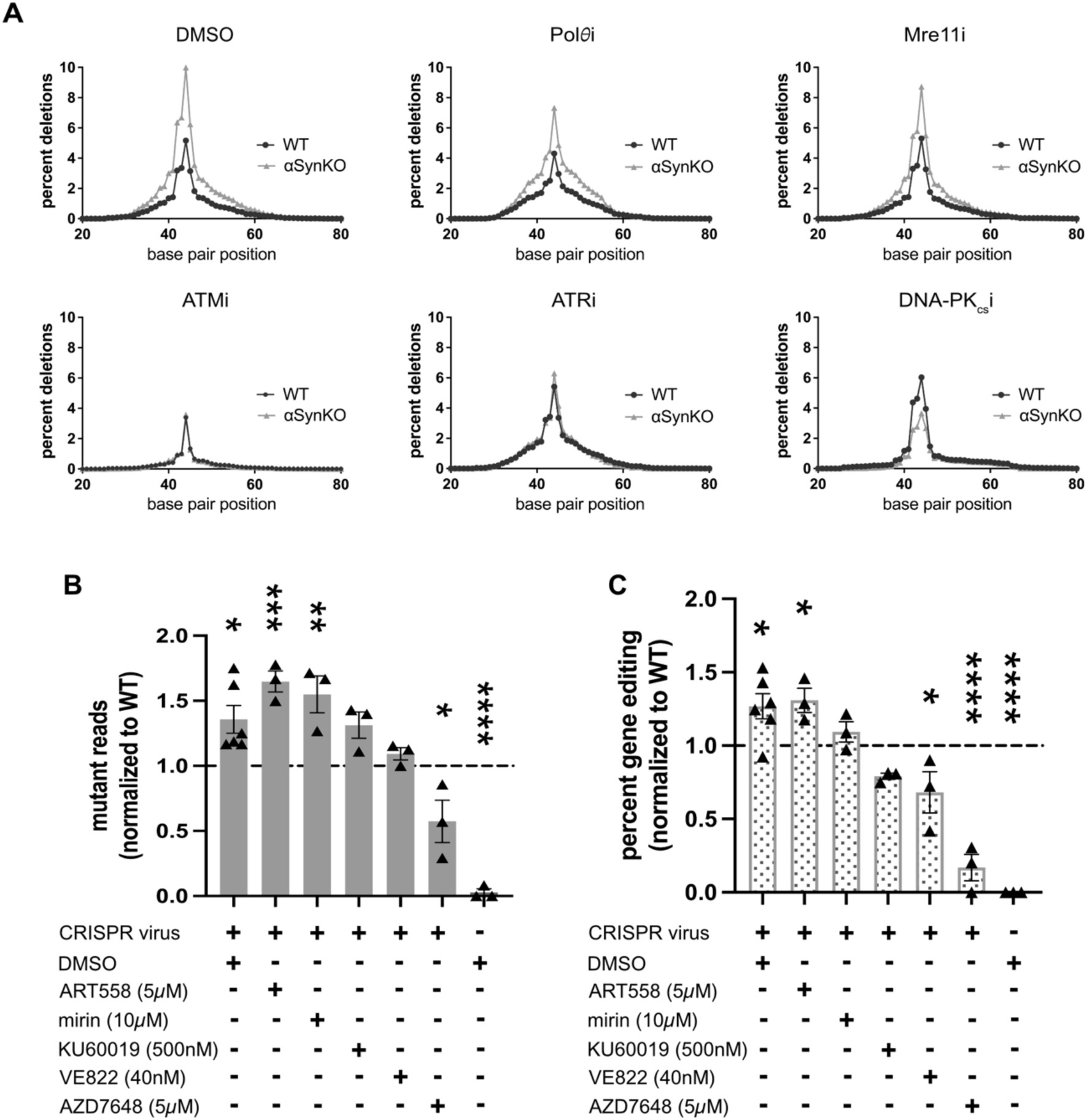
αSyn’s modulation of DSB repair in human cells is reversed by DNA-PK_cs_ inhibition. A) NGS of repair junction with percent of reads containing insertions (green) or deletions (blue) from WT and αSyn KO HAP1 cells transduced with CRISPR/Cas9 DSB inducing lentivirus. Mutant reads plotted against base pair position with cut site at base pair 45. Cells treated with 0.01% DMSO, Polθi (ART558 5μM), Mre11i (mirin 10μM), ATMi (KU60019 500nM), ATRi (VE822 40nM), DNA-PK_cs_i (AZD7648 5μM). B) Quantification of NGS of αSyn KO samples with condition normalized to WT DMSO. ANOVA summary p<0.0001. Post-hoc multiple comparisons: Significant increase of mutant reads in repair junction of DMSO treated αSyn KO cells (1.358 ±0.166) compared to DMSO treated WT cells (p=0.0124), in Polθi treated αSyn KO cells (1.648 ±0.081)(p=0.0002), and in Mre11i treated αSyn KO cells (1.549 ±0.142)(p=0.0014). ATMi treated αSyn KO cells (1.313 ±0.101)(p=0.1162). ATRi treated αSyn KO cells (1.093 ±0.047)(p=0.9816). Significant decrease in mutant reads of DNA-PK_cs_i treated αSyn KO cells (0.5734 ±0.163)(p=0.0154). αSyn KO Nontargeting Virus (0.028 ±0.028)(p<0.0001). N= 3 biological replicates, 1-2 technical replicate per biological replicate. One-way ANOVA. C) T7 Endonuclease I enzymatic assay quantification of percent gene editing of WT and αSyn KO cells treated with inhibitors from A) and C). % Gene Editing = 100 x (1 – (1-fraction cleaved)^1/2^). ANOVA summary p<0.0001. Post-hoc multiple comparisons: Significant increase of gene editing in DMSO treated αSyn cells (1.268 ±0.123) compared to DMSO treated WT cells (p=0.0212), and in Polθi treated αSyn KO cells (1.309 ±0.082)(p=0.0331). Mre11i treated αSyn KO cells (1.093 ±0.070)(p=0.9371). ATMi treated αSyn KO cells (0.7914 ±0.029)(p=0.2584). Significant decrease in mutant reads of ATRi treated αSyn KO cells (0.6812 ±0.106)(p=0.0262), and in DNA-PK_cs_i treated αSyn KO cells (0.1694 ±0.090)(p<0.0001). αSyn KO Nontargeting Virus (0.0 ±0.0)(p<0.0001). N= 3 biological replicates, 1-2 technical replicate per biological replicate. One-way ANOVA.

We also investigated the role of three phosphatidylinositol 3-OH-kinase-related kinases (PI3KKs) that are known to phosphorylate H2AX to produce γH2AX, ATM, ATM-and Rad3-related (ATR), and DNA-dependent protein kinase catalytic subunit (DNA-PK_cs_). We inhibited these three kinases with KU60019, VE822, and AZD7648, respectively. To determine the maximum concentration without toxic effects, we performed dose response curves for all 5 inhibitors in these cell lines and chose the maximum dose that did not cause toxicity (*data not shown*). Interestingly, we found a consistent effect in both our sequencing and T7EI assays that inhibiting DNA-PK_cs_ reversed the indel frequency effect normally seen in αSyn KO cells. In our previous experiments αSyn KO cells always had a higher frequency of indels created during the repair process. However, under the condition of DNA-PK_cs_ inhibition, this effect was reversed and αSyn KO cells had a lower deletion frequency than WT (Fig. 4B-C). As a control, we tested whether these inhibitors influenced cell proliferation and no significant changes were observed (Supplementary Fig. 1). These results suggest that αSyn’s normal function may be to modulate forms of c-NHEJ where DNA-PK_cs_ is important, such as those which require the DNA-PK_cs_-stimulated endonuclease activity of Artemis^55^, since our data is consistent with αSyn loss-of-function skewing repair towards a pathway that increases small indel frequency.

In order to assess whether αSyn re-expression is sufficient to reverse the abnormalities we detected in DSB repair in αSyn KO cells, we used transient transfection to express WT αSyn and a variety of mutants in WT and αSyn KO cells. We transfected WT αSyn, and 6 other mutant forms (A53T, E46K, A30P, delNAC 61-95 (deletion in residues 61-95), S129A, S129D) into WT and αSyn KO HAP1 cells and measured the indel frequency at the repair junction after using our CRISPR/Cas9 DSB induction system targeting the *DNMT3B* gene as previously described. We confirmed rescue of αSyn expression for WT and all mutant forms via western blot (Supplementary Fig. 2), but unfortunately, the transient transfection approach to re-express αSyn altered our system so that there was no difference between WT and αSyn KO cells in indel frequency as assayed by sequencing or the T7EI assay, as we had previously detected. This made it difficult to interpret the effect of our attempted re-expression in this context where we found no significant difference between transfected WT and αSyn KO cells in indel frequency (Supplementary Fig. 2) or in their proliferation (Supplementary Fig. 3).

### Inhibition of PLK protects against neurodegeneration in Lewy pathology mouse model *in vivo*

We next wanted to test how manipulating phosphorylation of αSyn may affect cell death in a Lewy pathology mouse model, given our recent work suggesting that pSyn binds DNA very differently than unphoshorylated αSyn^28^. We measured cell survival of Lewy inclusion-containing neurons longitudinally over 4 weeks utilizing an *in vivo* multiphoton imaging approach in our previously characterized A53T Syn-GFP mouse line^56^. Previous work shows that the GFP tag used does not affect synuclein aggregation in this experimental paradigm^57,58^. Mouse cortical regions were imaged for a 2 week baseline period, and then for an additional 2 weeks during exposure to the PLK inhibitor BI2536 or saline control. This PLK inhibition started at day 60 after αSyn preformed fibril (PFF) injection to induce Lewy pathology, a time point we have previously shown leads to robust cortical Lewy pathology that can be imaged *in vivo* with this approach^56^. No significant differences in the rate of Lewy inclusion-bearing neuron cell death were detected between the two groups of mice during the baseline imaging period. However, after the treatment period had begun, we measured an increase in survival rate of cells bearing Lewy inclusions in mice treated with twice-weekly BI2536 injections compared to those receiving saline control injections (Fig. 5A-5B). These results suggest that acute pharmacologic inhibition of PLK protects against neurodegeneration of neurons bearing Lewy inclusions and extends the result we obtained previously in PLK2 KO mice^50^ by reproducing this neuroprotective effect with a more clinically relevant treatment paradigm.

**Figure 5:**
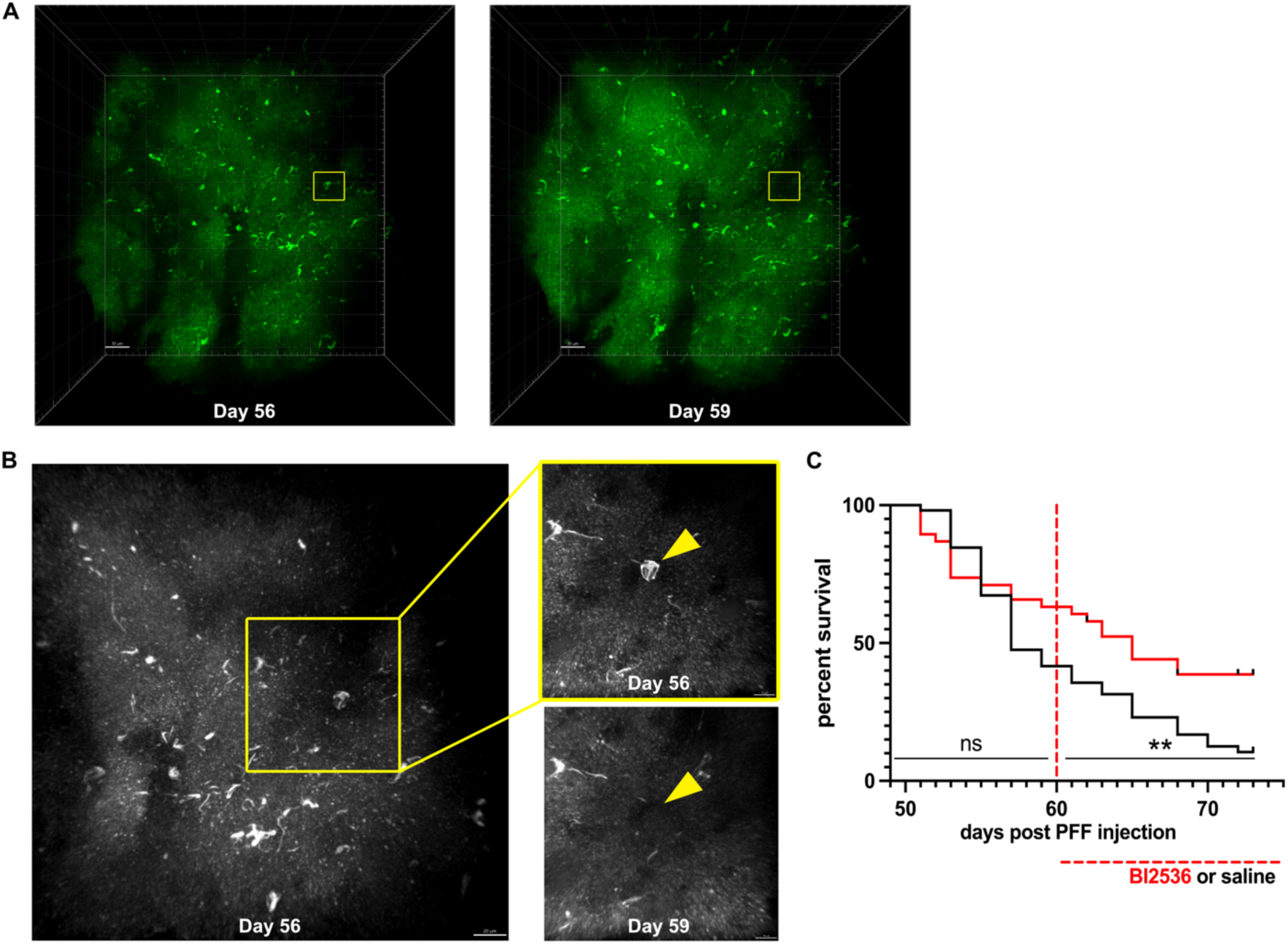
PLK inhibition protects against neurodegeneration in neurons bearing somatic Lewy inclusions *in vivo*. A) 3D Reconstruction of mouse brain cortical pathology *in vivo* on Day 55 and Day 59 post PFF injection with inclusion represented in B) highlighted (yellow square). Scale bar 30μm. 255μm z-stack. B) Representative images of mouse brain cortex *in vivo* demonstrating loss of cell body bearing αSyn somatic inclusion (yellow arrowhead) from day 55 to day 59 post PFF injection. Scale bar 20μm (LEFT), scale bar 10μm (RIGHT). Images were taken as separate acquisitions of inclusions (top 21μm z-stack, bottom 36μm z-stack). C) Survival curve of somatic inclusions across 25 days of longitudinal imaging of cortical regions *in vivo* in mice treated with saline or PLK 1/2/3/4 Inhibitor BI2536 (15mg/kg) IP injections for 2 weeks starting day 60 post PFF injection. Overall Mantel-cox test p=0.0161. Pre BI2536/saline treatment p=0.1454. Post BI2536/saline treatment p<0.0055. Saline treated group N= 4 animals. BI2536 treated group N=4 animals. 108 inclusions counted.

We next tested how this inhibition of PLK may affect αSyn’s role in DSB repair. We set out to do this by treating HAP1 cells with BI2536 to test its effect on αSyn modulated DSB repair in our CRISPR/Cas9-mediated DSB induction paradigm, but unfortunately, BI2536 treatment was toxic to HAP1 cells at the relevant concentrations. Because of this, we tested another PLK inhibitor GW843682X which had a better toxicity profile on HAP1 cells in our hands. We found an increased frequency of indels at the DSB repair junction of WT and αSyn KO cells when treated with GW843682X compared to vehicle treated cells (Supplementary Fig. 4A). In mouse cortical neurons, no significant differences observed (Supplementary Fig. 4B). These differences between our *in vivo* imaging results and our DSB repair assay in cultured cells may be due to the specificity differences in the inhibitors we were required to use. BI2536 inhibits PLK1, 2 and 3 with a similar IC50, while GW843682X is more selective from PLK1 and 3, and previous work suggests that PLK2 may be the most important family member for phosphorylating αSyn^44,48,50,59^.

### Inhibition of PLK increases levels of aggregated αSyn within somatic inclusions

To investigate the effects of PLK inhibition on aggregated αSyn within Lewy pathology, we used fixed tissue immunohistochemstry (IHC) to study neuronal somatic inclusions from mouse cortex after BI2536 or saline treatment after our *in vivo* imaging experiment (Fig. 5) had ended. Interestingly, BI2536 treatment reduced the ratio of pSyn/αSyn as we predicted, but for reasons different than we originally expected. BI2536 treatment had no significant effect on absolute pSyn levels measured, but it did increase the level of αSyn protein within Lewy inclusions. This was the cause of the decrease in the pSyn/αSyn ratio we detected. This suggests that at PLK1, 2 and 3 are not specific Lewy pathology kinases, but can have effects on the levels of αSyn protein within the inclusion (Fig 6A-6B). Previous work has suggested that PLK inhibition leads to a decrease in degradation of αSyn within aggregates^49^. Our data is consistent with this result. We next tested whether PLK inhibition leads to changes in γH2AX levels, via IHC. We found that BI2536 treatment caused an increase in γH2AX levels, both in cells with and without somatic Lewy inclusions (Fig. 6C). How PLK inhibition leads to an increase in γH2AX is not clear; it could be due to a specific effect on mediators of DSB repair like C-terminal binding protein (CtBP) interacting protein (CtIP), which are known to be phosphorylated by PLK^60,61^ or potentially through its effect causing increased aggregated αSyn levels within cells.

**Figure 6.**
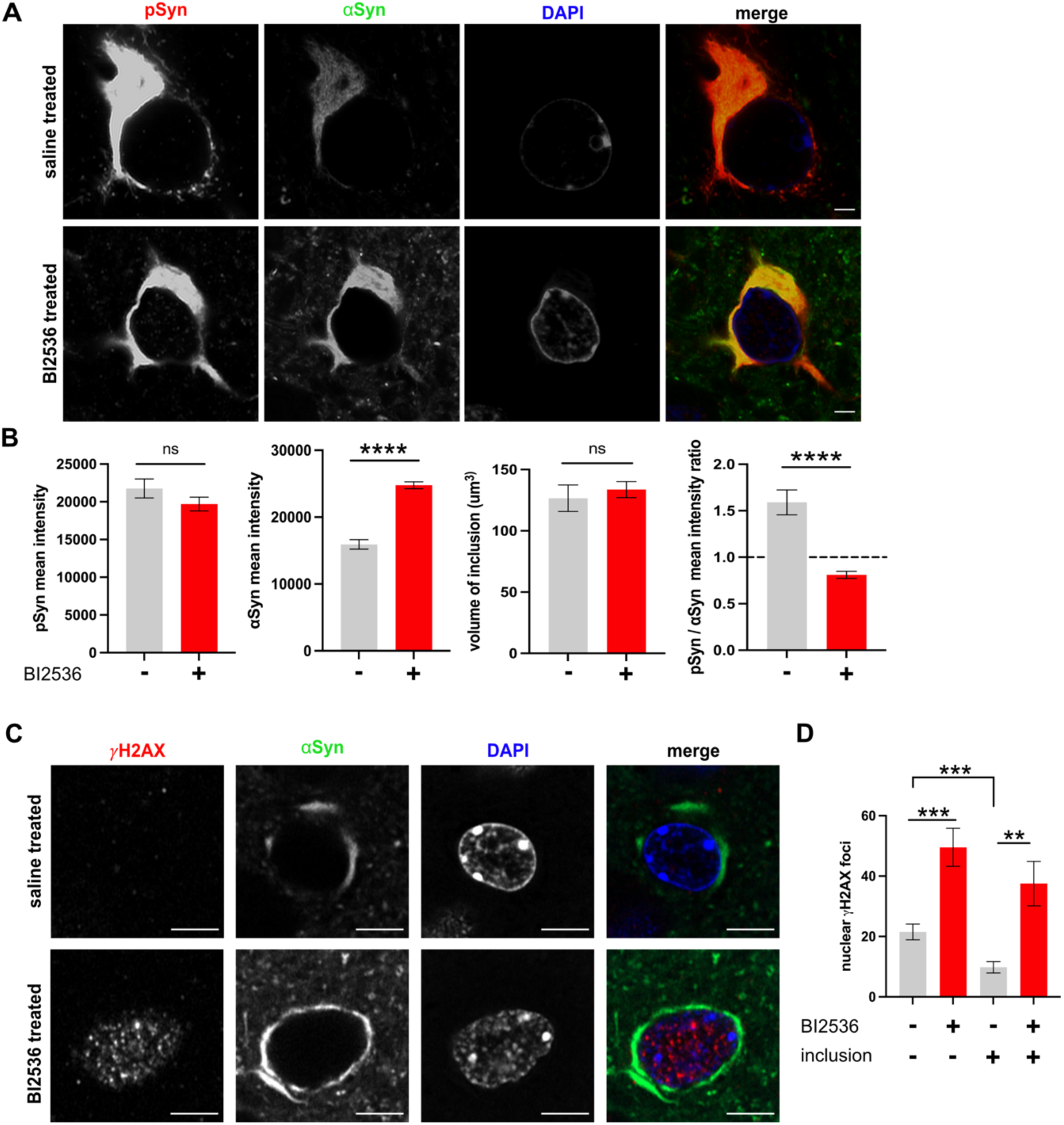
PLK inhibition increases levels of aggregated αSyn within somatic Lewy inclusions. A) BI2536 treatment (15mg/kg IP injections twice per week for two weeks) is associated with no change in pSyn levels, but increased total αSyn levels within PFF-induced aggregated somatic inclusion. Scale bar 2μm. B) Quantification of synuclein levels within the aggregate. No significant difference between pSyn mean intensity from saline treated mice (21766 ±1255.436) and BI2536 treated mice (19698 ±918.248)(p=0.1805). Significant increase of αSyn mean intensity from BI2536 treated mice (24755 ±511.682) compared to saline treated mice (15920 ±707.861)(p<0.0001). No significant difference of volume of the inclusion between saline treated mice (126.6 ±10.803) and BI2536 treated mice (133.6 ±6.606)(p=0.5559). Significant decrease of pSyn/αSyn mean intensity ratio of BI2536 treated mice (0.8106 ±0.038) compared to saline treated mice (1.590 ±0.135)(p<0.0001). N= 5-6 mice in each group, n=143 inclusions. Two-tailed student’s t-test. C) BI2536 treatment is associated with increased DSB levels in PFF-induced cortical Lewy pathology mouse model. Scale bar 5μm. RIGHT: Quantification. Significant increase of nuclear *γ*H2AX foci in cells without inclusions from BI2536 treated mice (49.55 ±6.334) compared to cells without inclusions from saline treated mice (21.54 ±2.605)(p=0.0007), but no significant difference when compared to cells bearing inclusions from BI2536 treated mice (37.53 ±7.357)(p=0.2178). Significant increase of nuclear *γ*H2AX foci in BI2536 treated cells bearing inclusions compared to saline treated cells bearing inclusions (9.812 ±1.886)(p=0.0031). Significant decrease of nuclear *γ*H2AX foci in saline treated cells bearing inclusions compared to cells without inclusions (p=0.0004). N= 5-6 mice in each group, n=349 inclusions. Two-tailed student’s t-test.

## Discussion

Here, we demonstrate that αSyn loss-of-function leads to impairment in the NHEJ form of DSB repair using a plasmid-based reporter assay in human cells (Fig. 1). Importantly, using a CRISPR/Cas9-based system to introduce a DSB within a single genomic location within a chromosomal context also showed an impairment in DSB repair in human cells (Fig. 2) and mouse cortical neurons (Fig. 3) in culture in the αSyn KO condition. The effect of αSyn KO is to increase the frequency of small indels found in the repaired DNA, suggesting that αSyn might function to promote forms of DSB repair that have a lower frequency of small indels, for example, sub-forms of c-NHEJ. Interestingly, DNA-PK_cs_ inhibition reversed this effect and led to lower indel levels in the αSyn KO condition (Fig. 4). Inhibition of PLK in our *in vivo* multiphoton imaging paradigm suggested that acutely inhibiting this kinase can improve the survival of these neurons (Fig. 5) and that this was associated with a reduction in pSyn/αSyn ratio within Lewy inclusions because of an increase in aggregated αSyn levels (Fig. 6).

Although a normal role for αSyn in DSB repair is a relatively new concept that we recently introduced to the field, several other groups have linked perturbations in αSyn with DNA damage. For example, both increased expression of αSyn or the PD-associated A30P mutation is correlated with down-regulation of genes involved in DNA repair, while only WT αSyn expression, and not the A30P form, induces DNA damage in dopaminergic neurons^62^. Other recent work shows that oxidized, misfolded αSyn can directly lead to DNA damage via an endonuclease activity, and that iron-dependent DNA breaks are associated with the triplication of the SNCA gene in a PD patient-derived IPSC line *in vitro*^63^. Notably, αSyn pathology induced by PFF injection causes a DNA damage response *in vivo* signaled by increased *γ*H2AX and 53BP1 foci^64^. Our work suggests that in addition to this effect of dysregulated and/or aggregated αSyn contributing to DNA damage, a normal physiologic function of αSyn could be to repair DSBs. Our previous work provided evidence for this by showing recruitment of αSyn to the sites of DNA damage in HAP1 cells and mouse cortex *in vivo*, and that αSyn loss-of-function led to higher DSB and *γ*H2AX levels, including after treatment with the chemical inducer of DSBs, bleomycin^1^. The current study significantly extends these results and again suggests a normal function for αSyn is to repair DSBs, but now when they are induced in a selective way within a single site in genomic DNA, both in human cells and mouse cortical neurons. In addition, we have found that αSyn plays an important role to facilitate sub-forms of c-NHEJ which result in a lower frequency of indels during the DSB repair process when αSyn is present at normal expression levels. It is interesting to consider how our data fits in with the previous work from other groups suggesting that αSyn aggregation leads to increased levels of DNA damage. We propose, given our data presented here, that αSyn facilitates c-NHEJ through a mechanism that is dependent on DNA-PK_cs_ activity, and that disrupting this process by overexpressing, mutating or aggregating αSyn leads to aberrant DSB repair. This could be due to a combination of factors, including shifting NHEJ towards a form that causes an increase in small indel frequency during DSB repair. In neurons, which are heavily dependent on NHEJ forms of DSB repair, this could lead to a progressive increase in mutation burden with time in Lewy Body-containing cells. We speculate that when a certain level of these small indels accumulate, it may trigger dysregulation of critical gene expression and neuroinflammatory pathways that lead to programmed cell death. It will be important in future studies to test whether there is evidence for this in human patient-derived tissue or model systems exhibiting Lewy pathology using single-cell approaches, since our model directly predicts an increased frequency of small indels within the genome of Lewy Body-containing neurons.

One of our more unexpected results was the evidence for interactions between αSyn and DNA-PK_cs_ in our DSB repair assays. There are multiple main pathways to repair a DSB, including c-NHEJ, alt-NHEJ, SSA and HR. How the cell decides which pathway to use is a critical question and previous work suggests that DNA-PK_cs_ is an important component in this decision. Because of its cellular abundance, DNA-PK_cs_ is thought to likely be an early repair factor binding DSBs^52^ and shunting cells towards forms of c-NHEJ that can introduce small indels^55^ and away from alt-NHEJ and HR^65^. Consistent with this, our experiment blocking an alt-NHEJ polymerase, Pol *θ*, showed no effect (Fig. 4). In addition to the role DNA-PK_cs_ has in inhibiting HR, which can be regulated by autophosphorylation^66^, DNA-PK_cs_ may also regulate a cell’s decision between sub-forms of c-NHEJ and alt-NHEJ^67,68^, although other data suggests that DNA-PK_cs_ phosphorylation of ATM^69^ or Ku^70^ facilitates this choice. C-NHEJ can proceed via a pathway independent of DNA-PK_cs_, involving Ku binding to XRCC4 and ligase 4 that does not introduce indels, or via other forms requiring DNA-PK_cs_ activity that activates the endonuclease Artemis and introduces small deletions at the repair site^55^. Our data suggests that αSyn promotes sub-forms of c-NHEJ which do not introduce small indels when DNA-PK_cs_ is not inhibited. Our result that DNA-PK_cs_ inhibition reverses this effect and αSyn switches to increasing the frequency of small indels suggests that these two molecules do potentially interact in potentially complicated ways. This interaction may be direct or indirect, and could also potentially involve modulation of liquid-liquid phase separation processes that are relevant to DNA repair and neurodegeneration^71^, and in which αSyn has been recently implicated^72^. One possibility is that αSyn could be acting like a partial agonist for DNA-PK_cs_ activity. In the presence of DNA-PK_cs_’s endogenous activator, Ku protein at DSB sites, αSyn acts to reduce the level of DNA-PK_cs_ kinase activity, thereby reducing Artemis endonuclease activity and reducing the level of small indels created at the repair junction. However, in the presence of strong DNA-PK_cs_ chemical inhibition, αSyn partial agonism could actually promote its kinase activity and promote Artemis-mediated endonuclease activity giving rise to small indel formation. Multiple models are possible, however, including ones where αSyn could directly promote or inhibit other DSB repair pathways that could influence the frequency of small indels formed during repair. It will be interesting in future experiments to test which of these models could be operative.

Our *in vivo* multiphoton imaging data suggests that inhibiting PLK acutely can improve survival of Lewy inclusion-bearing neurons in cortex. This extends our previous work in PLK2 KO mice showing a similar result^50^ by suggesting that pharmacologic inhibition may have similar effects and be potentially therapeutic. We originally expected that PLK inhibition with BI2536, which inhibits PLK1, 2 & 3, would have effects on pSyn levels within Lewy pathology, suggesting that PLK1 or 3 was a Lewy pathology kinase, since our previous work in PLK2 KO mice suggested that PLK2 was not a Lewy pathology kinase^50^. However, our fixed tissue IHC analysis did not suggest that pSyn levels were lower after BI2536 treatment, suggesting that PLK is not a Lewy pathology kinase *in vivo*. We did detect changes in total aggregated αSyn levels within inclusions, with PLK inhibition leading to an increase. Our data is consistent with previous work from Lashuel and colleagues that finds that PLK2 regulates and enhances autophagic clearance of αSyn in a kinase-dependent manner^49^. More investigation is required to decipher how PLK inhibition leads to increased neuronal survival and whether genomic stability plays a role, but it is interesting to speculate that increases in αSyn levels within the inclusion are mirrored by increases in potentially low levels of soluble nuclear αSyn as well that could be promoting more efficient DSB repair.

In summary, our data using powerful single genomic DSB induction approaches clearly demonstrates the importance of αSyn in NHEJ, favoring less error prone c-NHEJ pathways. This is the case both in a human cell line and in mouse primary cortical neurons, directly implicating αSyn-mediated DSB repair in this important cell type. Our data suggesting an interaction between αSyn and DNA-PK_cs_ in this process is the first time, to our knowledge, this has been suggested and sets the stage for future work to test whether αSyn could act as a partial agonist of DNA-PK_cs_ at DSBs. We also show how acute PLK inhibition can lead to neuroprotection in a Lewy pathology model and determining the mechanism for this effect could help lead to new treatments for clinically important forms of neurodegenerative disease.

## Methods and Materials

### Cell culture

HAP1 WT (item #C631 bath 29663) and HAP1 Human SNCA 103bp deletion knockout (item #HZGHC003210c003 batch 2) cell lines were purchased from Horizon Discovery. Cells were maintained at 37C 5% CO2 and grown in IMDM media (Gibco #11995-065)+10% Fetal Bovine Serum +5% penicillin-streptomycin (Gibco #1514022). Cells were passed when ∼75% confluent and discarded above passage number 10.

### Plasmid NHEJ and HR reporters

HAP1 WT and SNCA KO cells were seeded for ∼75% confluency and transfected with NHEJ and HR plasmid reporters^51^ using X-tremeGENE HP DNA Transfection Reagent (1:3 DNA:reagent) in OptiMEM (Gibco #31985062). Images were taken on a Zeiss Axio Observer.D1 outfitted with an Excelitas X-Cite 120 LED GFP light at 24, 48, 72 hours. After 48 hours cells were spun down and either submitted for flow cytometry or the DNA was processed with Qiagen’s QIAprep kit. DNA was digested with T5 exonuclease for 1 hour at 37C and cleaned up using Cytivia’s mag-bind beads. DNA was digested with fspI for 45 minutes at 37C and cleaned up again with mag-bind beads. DNA was normalized and sent to Azenta Life Sciences for AmpliconEZ illumina sequencing.

### Flow Cytometry

All experiments were completed with the help of OHSU’s Flow Cytometry core. GFP expression was measured in living cells, not fixed cells on the same day of experiment. HAP1 cells were trypsinized with 0.5% trypsin (Gibco #25300062) to transfer cells to an 0.6mL tube. Cells were incubated in trypsin for 10 minutes on ice. Cells were spun down and resuspended in PBS. Cells were spun down and resuspended in FACS Buffer (PBS +1% FBS). Cells were strained and submitted to OHSU’s Flow Cytometry for GFP expression analysis. Cells were first gated on a forward side scatter to exclude debris. Cells were next gated on forward side scatter height x forward side scatter area for doublet discrimination. GFP efficiency was measured by taking GFP positive singlets over the total amount of single cells. For CRISPR/Cas9 lentiviral DSB assays, there was no significant difference in GFP efficiency between HAP1 WT cells treated with GFP lentivirus (21.54 ±4.92) and HAP1 KO cells (14.88 ±4.29) p=0.3653. Mutation values were not adjusted.

### Incucyte proliferation

96 well image lock plates (Essen) were coated with 80μg/mL matrigel for 24 hours. 35K cells were plated per well in IMDM (10% FBS, 5% penicillin-streptomycin) with 6 replicates of WT and SNCA KO cells each and transferred to the Sartorius Incucyte S3 microplate holder maintained at 37C at 5% CO2. 4x brightfield scans were taken 3 hours after plating and continuously every 3 hours for the duration of the experiment. For proliferation assays post transfection, cells were seeded at 15K per well to account for the different time course of the experiment.

### CRISPR/Cas9 lentiviral double strand break repair assay

Human HAP1 WT and SNCA KO cells were maintained in IMDM (Gibco# 11995-065 +10% FBS, +5% penicillin-streptomycin) under passage number 10. Cells were seeded 16,000 cells per well in a 96 well plate coated in Poly-D-Lysine (Cultrex 343910001). 24 hours after seeding, cells were incubated with a Horizon Discovery Human DNMT3b mCMV-EGFP lentivirus (cat. #VSGH12131) (MOI 1) in 50uL of IMDM. 150uL of maintenance media was added to cells 5 hours after start of lentiviral treatment. Cells were harvested 72 hours after lentiviral treatment with 0.05% trypsin. Cells were lysed in 20% 5x Phusion HF Buffer, 5% 20mg/mL Proteinase K, and 5% 10mg/mL RNaseA and stored at -80C. The repair junction was amplified using a standard PCR reaction with Phusion HotStart. Human cell line repair junction 288bp product reference sequence: TTTCTGAGCACAGAGGGTACAGGCCGGCTCTTCTTCGAATTTTACCACCTGCTGAATTACTCACGCCCCAAGGAGGGTGATGACCGGCCGTTCTTCTGGATGTTTGAGAATGTTGTAGCCATGAAGGTTGGCGACAAGAGGGACATCTCACGGTTCCTGGAGGTGAGGGAATCTGGGGACCTGATTGTCACAGACAGCCAGGGCAGGGAAAGCGCTGCTGGCAGTGATGATTGGTGGGTGTTGCCAACATTGGGAATGACTTTCCCGTTCTTGGTCTGGCTAGATCCA with forward primer TTTCTGAGCACAGAGGGTACAG and reverse primer TGGATCTAGCCAGACCAAGAAC. Cut site at 45bp CC^CC. Samples underwent a PCR cleanup protocol according to Qiagen’s QIAquick PCR Purification Kit protocol and DNA levels were measured via Qubit dsDNA HS Assay Kit. Samples were normalized to 20ng/uL and sent to Azenta Life Sciences for Next Generation Sequencing (AmpliconEZ).

Mouse E18 cortical neurons were transduced with an analogous mouse dnmt3b mCMV-EGFP lentivirus (cat. #VSGM12147) (MOI 0.35) in maintenance media DIV 6. Cells were harvested 72 hours after lentiviral treatment with 0.05% trypsin and submitted to the same protocol as listed above. Mouse neurons repair junction 272bp product reference sequence: CTGTGCTGTTCCCATTACAGAGGGCACAGGAAGGCTCTTCTTCGAGTTTTACCACTTGCTGAATTATACCCGCCCCAAGGAGGGCGACAACCGTCCATTCTTCTGGATGTTCGAGAATGTTGTGGCCATGAAAGTGAATGACAAGAAAGACATCTCAAGATTCCTGGCAGTGAGTGGATTGTCAGGGAAACCTGGCAGGGAAGGCGCCACTAACACGGAGGGCTGAGAAAATTATTTCCTGCTCAGAGGAGGGTGTGGCTTAATCTGAGAAC with forward primer CTGTGCTGTTCCCATTACAGAG and reverse primer GTTCTCAGATTAAGCCACACCC. Cut site at 52bp CC^CC.

### T7 endonuclease I assay

The repair junction of the DNMT3b gene from human HAP1 cell lysates from the DSBR Assay was amplified with the same PCR protocol to create a 544bp full length product. Samples were then heated for 10 min at 95C and slowly cooled for at 1C per minute to room temperature. Samples were digested with T7 Endonuclease I for 25 minutes at 37C and run out on a 2% agarose gel at 100V for 2 hours. Human DNMT3b gene 544bp product reference sequence: TGAGAAGGAGCCACTTGCTTCTGGCCAAGTTACTGGCAGCATCAGGGGCCTGTTGGTGCTGCCTACGCTCCATAGTAAATCCTCAGCCCACAAGGGAAATACCCTAGTAAATAGTGCCCTGCTGCTGCCTGTGTCCCTGCTGTCATTCAGGTGGACATAGACTGGTAGGCATCACCCTGAACTGTCAGGAGGCCATTGGGAACCTGCTGGTCTCAGGGAATAAGGTGGGTTGGGCTGGAGGTTTCAAATGAACCCTGCGCTGTCATCTTTTCTGAGCACAGAGGGTACAGGCCGGCTCTTCTTCGAATTTTACCACCTGCTGAATTACTCACGCCCCAAGGAGGGTGATGACCGGCCGTTCTTCTGGATGTTTGAGAATGTTGTAGCCATGAAGGTTGGCGACAAGAGGGACATCTCACGGTTCCTGGAGGTGAGGGAATCTGGGGACCTGATTGTCACAGACAGCCAGGGCAGGGAAAGCGCTGCTGGCAGTGATGATTGGTGGGTGTTGCCAACATTGGGAATGACTTTCCCGTTCTTGGTCTGGCTAGAT CCA with forward primer TGAGAAGGAGCCACTTGCTT and reverse primer ACTGAAAGGGCAAGAACCAG. Mouse dnmt3b gene 561bp product reference sequence: ACTTGGTGATTGGTGGAAGCCCATGCAATGATCTCTCTAACGTCAATCCTGCCCGCAAAGGTTTATATGGTAAGCAGGGTTTGGGAACCTCCAGCACCACTATGTGCCATGTGTCTATGTTCAAATGGAAAATGGAGAAAAGAAGCTGTTGTCAGTTGTTCAGCCGTATTCATGACTCAGGCCCGGTCCTTCCCAGACACAACAAATCCAGTTGCTTTCTTTTACTGCAGTGTCCTGGGGACACTCTTGGTCTTTTGAGGCTCGTTTGGAATGAAGGCTTTGACTAAACCTTGTCCTCCTGTGCTGTTCCCATTACAGAGGGCACAGGAAGGCTCTTCTTCGAGTTTTACCACTTGCTGAATTATACCCGCCCCAAGGAGGGCGACAACCGTCCATTCTTCTGGATGTTCGAGAATGTTGTGGCCATGAAAGTGAATGACAAGAAAGACATCTCAAGATTCCTGGCAGTGAGTGGATTGTCAGGGAAACCTGGCAGGGAAGGCGCCACTAACACGGAGGGCTGAGAAAATTATTTCCTGCTCAGAGGAGGGTGTGGCTT AA with forward primer ACTTGGTGATTGGTGGAAGC and reverse primer GTCTCCTCCCACACCGAATT.

### Animals

All mice lines were housed in OHSU’s Department of Comparative Medicine (DCM) facilities in a light-dark cycle vivarium. Animals were maintained under ad libitum food and water diet. All animal protocols were approved by OHSU IACUC, and all experiments were performed with every effort to reduce animal lives and animal suffering.

### Transgenic mouse lines

WT (C57BL/6NJ Strain#: 005304) and SNCA KO (C57BL/6N-*Sncatm1Mjff*/J Strai #: 016123) were obtained from The Jackson Laboratory. For embryonic cortical dissections, timed pregnancies were performed by breeding SNCA KO mice together or WT mice together for 72 hours and separating the female.

The A53T-Syn-GFP mouse line was genetically created^1^ and characterized^56^ according to our previous research.

### Primary Cultured Neurons

Cortices were dissected from E18 mice and kept in Hibernate-E (Gibco #A1247601) at 4C until dissociation within 1 to 2 hours. Primary cortical neurons were cultured according to an adapted protocol from the Banker Lab^73^. Cortices were transferred to HBSS (Gibco #14025092) on ice. Neurons were dissociated with 2.5% trypsin (Gibco #15090046) for 15 minutes at 37C, gently inverting every 5 minutes. Neurons were dissociated in sterile filtered plating medium (Neurobasal Medium Gibco #21103049 +10% FBS, +1% GlutaMAX Supplement, +1% sodium pyruvate, +1% penicillin-streptomycin) and frozen down with MD Bioproduct’s NeuroFreeze kit. Neurons were kept in liquid nitrogen up to 4 months. Neurons were thawed according to the NeuroFreeze kit and plated for 4 hours in PDL coated 96 well plates. Media was exchanged for sterile filtered maintenance media (Neurobasal Medium +2% B-27 Supplement 50X Gibco #17504044, +0.5% GlutaMAX Supplement) and neurons remained in a humidified incubator at 37C with 5%CO2 until DIV 5. One third of the media was exchanged for fresh maintenance media on DIV 5. Lentiviral Transduction for the DSB repair assay occurred on DIV 6.

### Pharmacological Agents

All inhibitors were purchased, stored at -20C, diluted in DMSO, aliquoted, and immediately stored at -80C. Treatment concentrations were decided by a treatment curve, selecting the highest concentration possible without affecting cell health. Final concentrations were between 2x-2000x each inhibitor’s IC50 for each desired target. 1000x Inhibitors were thawed and added to IMDM at 0.1%. Cells were treated with inhibitors concurrently with lentiviral treatment. Final Concentrations: 40nM ATRi (VE822), 500nM ATMi (KU60019), 200nM DNAPKi (NU7441), 5μM DNAPKi (AZD7648), 5μM Pol *θ*i (ART558), and 10μM Mre11i (mirin).

### Transfection

The mammalian expression vector pBApo-CMV-Pur (cat. #3421) was purchased from Takara. We used Genescript to insert the 144 amino acid sequence for WT synuclein into the backbone and create 6 mutated strains: S129A, S129D, A53T, E46K, A30P, and delNAC(61-95). Plasmids were then transformed, miniprepped, and stored at -20C. 24 hours before transfection, HAP1 cells were seeded to be 60% confluent. The jetOPTIMUS reagent was added to the DNA+jetOPTIMUS buffer according to the Polyplus protocol using 0.15ug of DNA for each well in a 96 well format, 1:1.5 ratio DNA : jetOPTIMUS reagent. Transfection was performed in IMDM +5% FBS for 4 hours and then media was exchanged for IMDM +10% FBS, +5% penicillin-streptomycin. Reagents were scaled up for transfections in a 60mm dish for subsequent western blot confirmation.

### Western Blot

The western blot protocol used in the paper was completed in accordance to previously published work^1^. Primary antibodies used were: anti-Syn 4B12, 1:500 dilution, mouse monoclonal, Biolegend, cat. 807804; GAPDH, 1:10,000 dilution, mouse monoclonal, Millipore, cat. MAB374.

### Mouse brain *in vivo* imaging & analysis

2 to 3 month-old male and female mice were injected with mouse WT sequence PFFs using the same protocol as we have previously published^1^. Cranial window surgeries were performed 5 weeks post PFF injection according to our previous published protocols. Mouse cortex was imaged 3 weeks post cranial window surgery using a Zeiss LSM 7MP multiphoton microscope. Zeiss Zen image acquisition software was used to collect z-stacks from layer 1 to layer 2/3 of the cortex with 3μm intervals at 63x zoom 1. Regions of interest (ROIs) were analyzed in FIJI and inclusions were verified visually for each day of imaging by hand. New inclusions were counted for each day of imaging and scored by hand. No detectible sex differences were observed. Survival curves were created with Prism10 (GraphPad). Cortical regions were imaged for 4 weeks at 3 times per week. After 2 weeks animals were given a 2 week treatment of saline or 15mg/kg BI2536 IP injections twice per week. Animals were sacrificed after 4 weeks of imaging for IHC.

### Immunofluorescence and Immunohistochemistry

#### HAP1 cells

Cells were seeded on #1.5 coverslips or glass-bottomed 96 well plates coated in PLL. Cells were fixed, permeabilized, incubated in blocking buffer, stained with primary and secondary antibodies, and mounted according to previously published protocols^1^. Cells were imaged on a Zeiss 980 laser-scanning confocal microscope with Zen software. Z-stacks of optimal intervals were acquired at 63x zoom 1.

#### Primary Cortical Neurons

Neurons were seeded into PDL coated Cellvis #1.5H glass-bottomed 96 well plates. After 72 hour lentiviral treatment, neurons were fixed on DIV 9 with 4% PFA for 15 minutes and permeabilized with 0.25% Triton X-100 for 20 minutes at RT. Neurons were incubated with Biotium NucSpot555/570 (cat. 41033) for 10 minutes at RT. Whole well images were acquired same day on a Zeiss Celldiscoverer 7 at 20x zoom 1. Images were stitched together using Zen software and an Imaris machine learning analysis protocol was used to exclude dead nuclei. A mean intensity 488 threshold was used to count GFP positive cells on Imaris and GFP efficiency was calculated. For CRISPR/Cas9 lentiviral DSB assays, there was no significant difference in GFP efficiency between WT E18 cortical neurons treated with GFP lentivirus (6.267 ±0.70) and KO E18 cortical neurons (16.05 ±6.18) p=0.1907. Mutation values were not adjusted.

#### Mouse Tissue

Brains from 4-5 month old mice were dissected and fixed according to previously published protocols^1^. Brains were sectioned into 50μM coronal slices using a Vibratome LeicaVT1000S. Tissue was fixed, permeabilized, incubated in blocking buffer, stained with primary and secondary antibodies, and mounted as previously published^1^. Imaging was performed on a Zeiss 980 laser-scanning confocal microscope with a 63x oil objective zoom 7.0. Z-stacks of the αSyn inclusion and nucleus with optimal intervals. Analysis was performed in IMARIS using a 3D surface reconstruction of the inclusion and the DAPI channel to create a nuclear mask. We then used IMARIS to quantify the nuclear *γ*H2AX foci count or αSyn and pSyn levels with an average intensity measurement.

Primary antibodies used were: anti-Syn1, 1:500 dilution, mouse monoclonal, BD Biosciences, cat. 610786; anti-Phospho-Histone H2a.X, 1:500 dilution, rabbit monoclonal, Cell Signaling, cat. 9718; anti-PhosphoS129-Syn EPY1536Y, 1:500 dilution, rabbit monoclonal, Abcam ab51253. Secondary antibodies used were: Alexa Fluor 555 goat anti-mouse, Abcam ab150114; Alexa Fluor 647 donkey anti-rabbit, Jackson ImmunoResearch Laboratories 711605152.

## Acknowledgments

We would like to thank the lab of Kelvin Luk for providing ⍺syn PFFs, Lev Federov and the OHSU Transgenic Mouse Model Core for generation of A53T Syn-GFP mice, and the OHSU Advanced Light Microscopy core for their guidance and equipment on several experiments. We also thank Michael Cohen, Amanda McCullough, Ian Martin, and Peter McKinnon for helpful discussions. Lastly, we recognize Unni Lab members who helped with the revision process: Moriah Arnold, Dr. Sydney Boutros, Anna Bowman, Jessica Keating, Dr. Carlos Soto Faguas, and Elias Wisdom. This work was supported in part by the NIH (T32AG055378, R01NS102227), the David Johnson Family Foundation, and the Lacroute Fellows Program.

## Author Contributions

Conceptualization, EPR & VKU. Methodology, EPR, VRO, VG, VKU. Formal analysis, EPR, VRO, JSB, VKU. Investigation, EPR, VRO, VKU. Writing - original draft, EPR. Writing - review & editing, EPR, VRO, VG, VKU, Visualization - EPR & VKU. Supervision, VKU. Project administration, VKU. Funding acquisition, EPR & VKU.

## Data Availability

All materials, data, and protocols will be made available to readers without undue qualifications.

## References

1. Schaser, A. J. et al. Alpha-synuclein is a DNA binding protein that modulates DNA repair with implications for Lewy body disorders. Scientific Reports 9, 10919 (2019).

2. Greten-Harrison, B. et al. αβγ-Synuclein triple knockout mice reveal age-dependent neuronal dysfunction. PNAS 107, 19573–19578 (2010).

3. Steidl, J. V., Gomez-Isla, T., Mariash, A., Ashe, K. H. & Boland, L. M. Altered short-term hippocampal synaptic plasticity in mutant alpha-synuclein transgenic mice. Neuroreport 14, 219–223 (2003).

4. Parihar, M. S., Parihar, A., Fujita, M., Hashimoto, M. & Ghafourifar, P. Alpha-synuclein overexpression and aggregation exacerbates impairment of mitochondrial functions by augmenting oxidative stress in human neuroblastoma cells. The International Journal of Biochemistry & Cell Biology 41, 2015–2024 (2009).

5. Kumar, V. et al. Alpha-synuclein aggregation, Ubiquitin proteasome system impairment, and l-Dopa response in zinc-induced Parkinsonism: resemblance to sporadic Parkinson’s disease. Mol Cell Biochem 444, 149–160 (2018).

6. McNaught, K. S. P., Belizaire, R., Isacson, O., Jenner, P. & Olanow, C. W. Altered proteasomal function in sporadic Parkinson’s disease. Exp Neurol 179, 38–46 (2003).

7. Cuervo, A. M., Stefanis, L., Fredenburg, R., Lansbury, P. T. & Sulzer, D. Impaired degradation of mutant alpha-synuclein by chaperone-mediated autophagy. Science 305, 1292–1295 (2004).

8. Winslow, A. R. et al. α-Synuclein impairs macroautophagy: implications for Parkinson’s disease. J Cell Biol 190, 1023–1037 (2010).

9. Xilouri, M., Brekk, O. R. & Stefanis, L. Autophagy and Alpha-Synuclein: Relevance to Parkinson’s Disease and Related Synucleopathies. Mov Disord 31, 178–192 (2016).

10. Maroteaux, L., Campanelli, J. & Scheller, R. Synuclein: a neuron-specific protein localized to the nucleus and presynaptic nerve terminal. J Neurosci 8, 2804–2815 (1988).

11. Burré, J., Sharma, M. & Südhof, T. C. Cell Biology and Pathophysiology of α-Synuclein. Cold Spring Harb Perspect Med 8, a024091 (2018).

12. Wang, L. et al. α-synuclein multimers cluster synaptic vesicles and alenuate recycling. Curr Biol 24, 2319–2326 (2014).

13. Zharikov, A. D. et al. shRNA targeting α-synuclein prevents neurodegeneration in a Parkinson’s disease model. J Clin Invest 125, 2721–2735 (2015).

14. Benskey, M. J. et al. Silencing Alpha Synuclein in Mature Nigral Neurons Results in Rapid Neuroinflammation and Subsequent Toxicity. Front Mol Neurosci 11, 36 (2018).

15. Goers, J. et al. Nuclear localization of alpha-synuclein and its interaction with histones. Biochemistry 42, 8465–8471 (2003).

16. Pinho, R. et al. Nuclear localization and phosphorylation modulate pathological effects of alpha-synuclein. Human Molecular Genetics 28, 31–50 (2019).

17. Kontopoulos, E., Parvin, J. D. & Feany, M. B. α-synuclein acts in the nucleus to inhibit histone acetylation and promote neurotoxicity. Human Molecular Genetics 15, 3012–3023 (2006).

18. Geertsma, H. M. et al. Constitutive nuclear accumulation of endogenous alpha-synuclein in mice causes motor impairment and cortical dysfunction, independent of protein aggregation. Hum Mol Genet 31, 3613–3628 (2022).

19. Lee Clough, R. & Stefanis, L. A novel pathway for transcriptional regulation of α-synuclein. The FASEB Journal 21, 596–607 (2007).

20. Somayaji, M., Lanseur, Z., Choi, S. J., Sulzer, D. & Mosharov, E. V. Roles for α-Synuclein in Gene Expression. Genes 12, 1166 (2021).

21. Iwata, A., Miura, S., Kanazawa, I., Sawada, M. & Nukina, N. α-Synuclein forms a complex with transcription factor Elk-1. Journal of Neurochemistry 77, 239–252 (2001).

22. Hallacli, E. et al. The Parkinson’s disease protein alpha-synuclein is a modulator of processing bodies and mRNA stability. Cell 185, 2035–2056.e33 (2022).

23. Agamanolis, D. P. & Greenstein, J. I. Ataxia-Telangiectasia: Report of a Case With Lewy Bodies and Vascular Abnormalities within Cerebral Tissue. Journal of Neuropathology & Experimental Neurology 38, 475–489 (1979).

24. Eilam, R., Peter, Y., Groner, Y. & Segal, M. Late degeneration of nigro-striatal neurons in ATM−/− mice. Neuroscience 121, 83–98 (2003).

25. Huang, C.-H., Mirabelli, C. K., Jan, Y. & Crooke, S. T. Single-strand and double-strand deoxyribonucleic acid breaks produced by several bleomycin analogs. Biochemistry 20, 233– 238 (1981).

26. Caporossi, D., Ciafrè, S. A., Pilaluga, M., Savini, I. & Farace, M. G. Cellular responses to H2O2 and bleomycin-induced oxidative stress in L6C5 rat myoblasts. Free Radical Biology and Medicine 35, 1355–1364 (2003).

27. Mah, L.-J., El-Osta, A. & Karagiannis, T. C. γH2AX as a molecular marker of aging and disease. Epigenetics 5, 129–136 (2010).

28. Dent, S. E. et al. Phosphorylation of the aggregate-forming protein alpha-synuclein on serine-129 inhibits its DNA-bending properties. Journal of Biological Chemistry 298, (2022).

29. Anderson, J. P. et al. Phosphorylation of Ser-129 Is the Dominant Pathological Modification of α-Synuclein in Familial and Sporadic Lewy Body Disease. J. Biol. Chem. 281, 29739–29752 (2006).

30. Tenreiro, S., Eckermann, K. & Outeiro, T. F. Protein phosphorylation in neurodegeneration: friend or foe? Front Mol Neurosci 7, 42 (2014).

31. Paleologou, K. E. et al. Phosphorylation at Ser-129 but not the phosphomimics S129E/D inhibits the fibrillation of alpha-synuclein. J. Biol. Chem. 283, 16895–16905 (2008).

32. Fujiwara, H. et al. alpha-Synuclein is phosphorylated in synucleinopathy lesions. Nat. Cell Biol. 4, 160–164 (2002).

33. Samuel, F. et al. Effects of Serine 129 Phosphorylation on α-Synuclein Aggregation, Membrane Association, and Internalization. J. Biol. Chem. 291, 4374–4385 (2016).

34. Ma, M.-R., Hu, Z.-W., Zhao, Y.-F., Chen, Y.-X. & Li, Y.-M. Phosphorylation induces distinct alpha-synuclein strain formation. Sci Rep 6, 37130 (2016).

35. Chen, L. & Feany, M. B. α-Synuclein phosphorylation controls neurotoxicity and inclusion formation in a Drosophila model of Parkinson disease. Nature Neuroscience 8, 657–663 (2005).

36. McFarland, N. R. et al. Alpha-Synuclein S129 Phosphorylation Mutants Do Not Alter Nigrostriatal Toxicity in a Rat Model of Parkinson Disease. J Neuropathol Exp Neurol 68, 515–524 (2009).

37. Gorbatyuk, O. S. et al. The phosphorylation state of Ser-129 in human α-synuclein determines neurodegeneration in a rat model of Parkinson disease. Proc Natl Acad Sci U S A 105, 763–768 (2008).

38. Kawahata, I., Finkelstein, D. I. & Fukunaga, K. Pathogenic Impact of α-Synuclein Phosphorylation and Its Kinases in α-Synucleinopathies. Int J Mol Sci 23, 6216 (2022).

39. Pronin, A. N., Morris, A. J., Surguchov, A. & Benovic, J. L. Synucleins are a novel class of substrates for G protein-coupled receptor kinases. J. Biol. Chem. 275, 26515–26522 (2000).

40. Wu, W., Sung, C. C., Yu, P., Li, J. & Chung, K. K. K. S-Nitrosylation of G protein-coupled receptor kinase 6 and Casein kinase 2 alpha modulates their kinase activity toward alpha-synuclein phosphorylation in an animal model of Parkinson’s disease. PLoS One 15, e0232019 (2020).

41. Ishii, A. et al. Casein kinase 2 is the major enzyme in brain that phosphorylates Ser129 of human alpha-synuclein: Implication for alpha-synucleinopathies. FEBS LeN 581, 4711–4717 (2007).

42. Qing, H., Wong, W., McGeer, E. G. & McGeer, P. L. Lrrk2 phosphorylates alpha synuclein at serine 129: Parkinson disease implications. Biochem. Biophys. Res. Commun. 387, 149–152 (2009).

43. Basso, E. et al. PLK2 modulates α-synuclein aggregation in yeast and mammalian cells. Mol Neurobiol 48, 854–862 (2013).

44. Bergeron, M. et al. In vivo modulation of polo-like kinases supports a key role for PLK2 in Ser129 α-synuclein phosphorylation in mouse brain. Neuroscience 256, 72–82 (2014).

45. Aubele, D. L. et al. Selective and brain-permeable polo-like kinase-2 (Plk-2) inhibitors that reduce α-synuclein phosphorylation in rat brain. ChemMedChem 8, 1295–1313 (2013).

46. Waxman, E. A. & Giasson, B. I. Characterization of kinases involved in the phosphorylation of aggregated α-synuclein. J Neurosci Res 89, 231–247 (2011).

47. Mbefo, M. K. et al. Phosphorylation of Synucleins by Members of the Polo-like Kinase Family. J Biol Chem 285, 2807–2822 (2010).

48. Inglis, K. J. et al. Polo-like kinase 2 (PLK2) phosphorylates alpha-synuclein at serine 129 in central nervous system. J Biol Chem 284, 2598–2602 (2009).

49. OueslaG, A., Schneider, B. L., Aebischer, P. & Lashuel, H. A. Polo-like kinase 2 regulates selective autophagic α-synuclein clearance and suppresses its toxicity in vivo. Proc. Natl. Acad. Sci. U.S.A. 110, E3945–3954 (2013).

50. Weston, L. J. et al. Genetic deletion of Polo-like kinase 2 reduces alpha-synuclein serine-129 phosphorylation in presynaptic terminals but not Lewy bodies. Journal of Biological Chemistry 296, 100273 (2021).

51. Seluanov, A., Mao, Z. & Gorbunova, V. Analysis of DNA Double-strand Break (DSB) Repair in Mammalian Cells. J Vis Exp (2010) doi:10.3791/2002.

52. Neal, J. A. & Meek, K. Choosing the right path: does DNA-PK help make the decision? Mutat Res 711, 73–86 (2011).

53. Iliakis, G. et al. Mechanisms of DNA double strand break repair and chromosome aberration formation. Cytogenet Genome Res 104, 14–20 (2004).

54. Iliakis, G., Murmann, T. & Soni, A. Alternative end-joining repair pathways are the ultimate backup for abrogated classical non-homologous end-joining and homologous recombination repair: Implications for the formation of chromosome translocations. Mutat Res Genet Toxicol Environ Mutagen 793, 166–175 (2015).

55. Chang, H. H. Y., Pannunzio, N. R., Adachi, N. & Lieber, M. R. Non-homologous DNA end joining and alternative pathways to double-strand break repair. Nat Rev Mol Cell Biol 18, 495–506 (2017).

56. Schaser, A. J. et al. Trans-synaptic and retrograde axonal spread of Lewy pathology following pre-formed fibril injection in an in vivo A53T alpha-synuclein mouse model of synucleinopathy. Acta Neuropathologica Communications 8, 150 (2020).

57. Spinelli, K. J. et al. presynaptic Alpha-Synuclein Aggregation in a Mouse Model of Parkinson’s Disease. J Neurosci 34, 2037–2050 (2014).

58. Osterberg, V. R. et al. Progressive aggregation of alpha-synuclein and selective degeneration of lewy inclusion-bearing neurons in a mouse model of parkinsonism. Cell Rep 10, 1252– 1260 (2015).

59. Lou, H. et al. Serine 129 phosphorylation reduces the ability of alpha-synuclein to regulate tyrosine hydroxylase and protein phosphatase 2A in vitro and in vivo. J Biol Chem 285, 17648–17661 (2010).

60. Barton, O. et al. Polo-like kinase 3 regulates CtIP during DNA double-strand break repair in G1. Journal of Cell Biology 206, 877–894 (2014).

61. Wang, H. et al. PLK1 targets CtIP to promote microhomology-mediated end joining. Nucleic Acids Research 46, 10724–10739 (2018).

62. Paiva, I. et al. Sodium butyrate rescues dopaminergic cells from alpha-synuclein-induced transcriptional deregulation and DNA damage. Hum Mol Genet 26, 2231–2246 (2017).

63. Vasquez, V. et al. Chromatin-Bound Oxidized α-Synuclein Causes Strand Breaks in Neuronal Genomes in in vitro Models of Parkinson’s Disease. J Alzheimers Dis 60, S133–S150 (2017).

64. Milanese, C. et al. Activation of the DNA damage response in vivo in synucleinopathy models of Parkinson’s disease. Cell Death Dis 9, 818 (2018).

65. Mao, Z., Bozzella, M., Seluanov, A. & Gorbunova, V. Comparison of nonhomologous end joining and homologous recombination in human cells. DNA Repair (Amst) 7, 1765–1771 (2008).

66. Neal, J. A. et al. Inhibition of homologous recombination by DNA-dependent protein kinase requires kinase activity, is Gtratable, and is modulated by autophosphorylation. Mol Cell Biol 31, 1719–1733 (2011).

67. Udayakumar, D., Bladen, C. L., Hudson, F. Z. & Dynan, W. S. Distinct pathways of nonhomologous end joining that are differentially regulated by DNA-dependent protein kinase-mediated phosphorylation. J Biol Chem 278, 41631–41635 (2003).

68. Perrault, R., Wang, H., Wang, M., Rosidi, B. & Iliakis, G. Backup pathways of NHEJ are suppressed by DNA-PK. J Cell Biochem 92, 781–794 (2004).

69. Zhou, Y. et al. Regulation of the DNA Damage Response by DNA-PKcs Inhibitory Phosphorylation of ATM. Mol Cell 65, 91–104 (2017).

70. Falah, F. et al. Ku regulates the non-homologous end joining pathway choice of DNA double-strand break repair in human somatic cells. PLoS Genet 6, e1000855 (2010).

71. Webber, C. J., Lei, S. (Eric) & Wolozin, B. The pathophysiology of neurodegenerative disease: Disturbing the balance between phase separation and irreversible aggregation. Prog Mol Biol Transl Sci 174, 187–223 (2020).

72. Ray, S. et al. α-Synuclein aggregation nucleates through liquid–liquid phase separation. Nat. Chem. 12, 705–716 (2020).

73. Kaech, S. & Banker, G. Culturing hippocampal neurons. Nat Protoc 1, 2406–2415 (2006).

