## Supplemental Figures for "Alpha-synuclein regulates the repair of genomic DNA double-strand breaks in a DNA-PK_cs_-dependent manner"

*Supplemental Materials:*

**Supplemental Figure 1.**

**A**

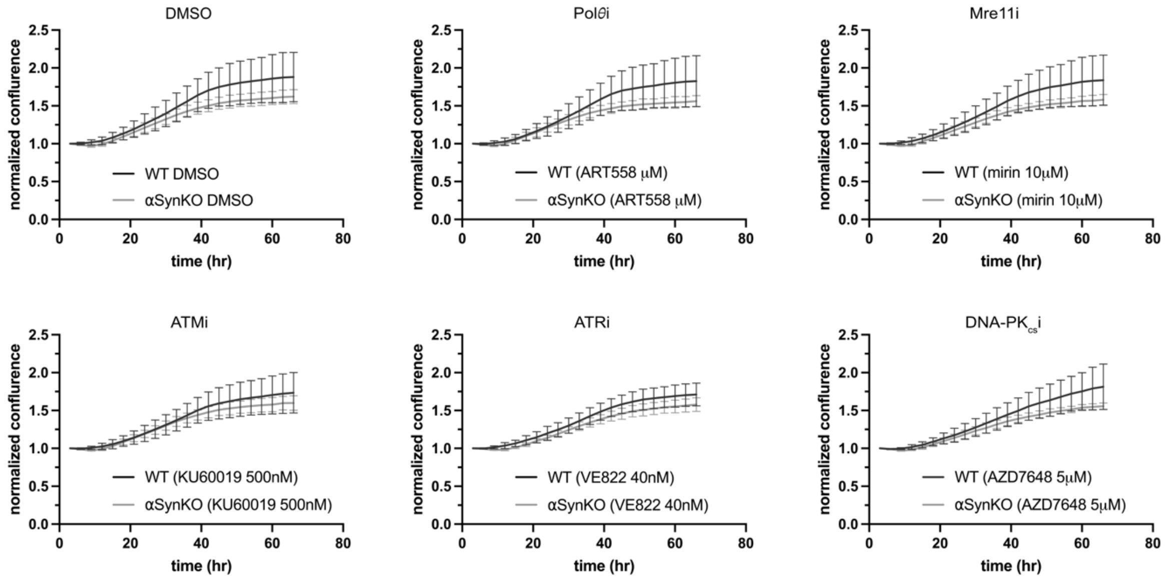


**Supplemental Figure 1. WT and αSyn KO HAP1 cell proliferation is unchanged by inhibition of Polθ/Mre11/ATM/ATR/DNA-PK$_{cs}$.**

A) Normalized confluence of WT and αSyn KO cells treated with 0.01% DMSO, Polθi (ART558 5uM), Mre11i (mirin 10μM), ATMi (KU60019 500nM), ATRi (VE822 40nM), DNA-PK$_{cs}$i (AZD7648 5μM). No significant differences observed. WT DMSO= 95% CI 50% maximum (27.25-61.40). KO DMSO= 95% CI 50% maximum (27.11-32.87). WT Polθi= 95% CI 50% maximum (27.46-69.68). KO Polθi= 95% CI 50% maximum (25.87-30.66). WT Mre11i= 95% CI 50% maximum (28.76-117.6). KO Mre11i= 95% CI 50% maximum (28.85-34.01). WT ATMi=95% CI 50% maximum (28.52-75.11). KO ATMi= 95% CI 50% maximum (27.98-33.98). WT ATRi= 95% CI 50% maximum (30.55-46.05). KO ATRi= 95% CI 50% maximum (31.14-37.96). WT DNA-PK$_{cs}$i = 95% CI 50% maximum (34.08-poor fit). KO DNA-PK$_{cs}$i = 95% CI 50% maximum (32.79-40.85). N=3 biological replicates, 6 technical replicates per biological replicate). Sigmoidal nonlinear regression.

**Supplementary Figure 2.**

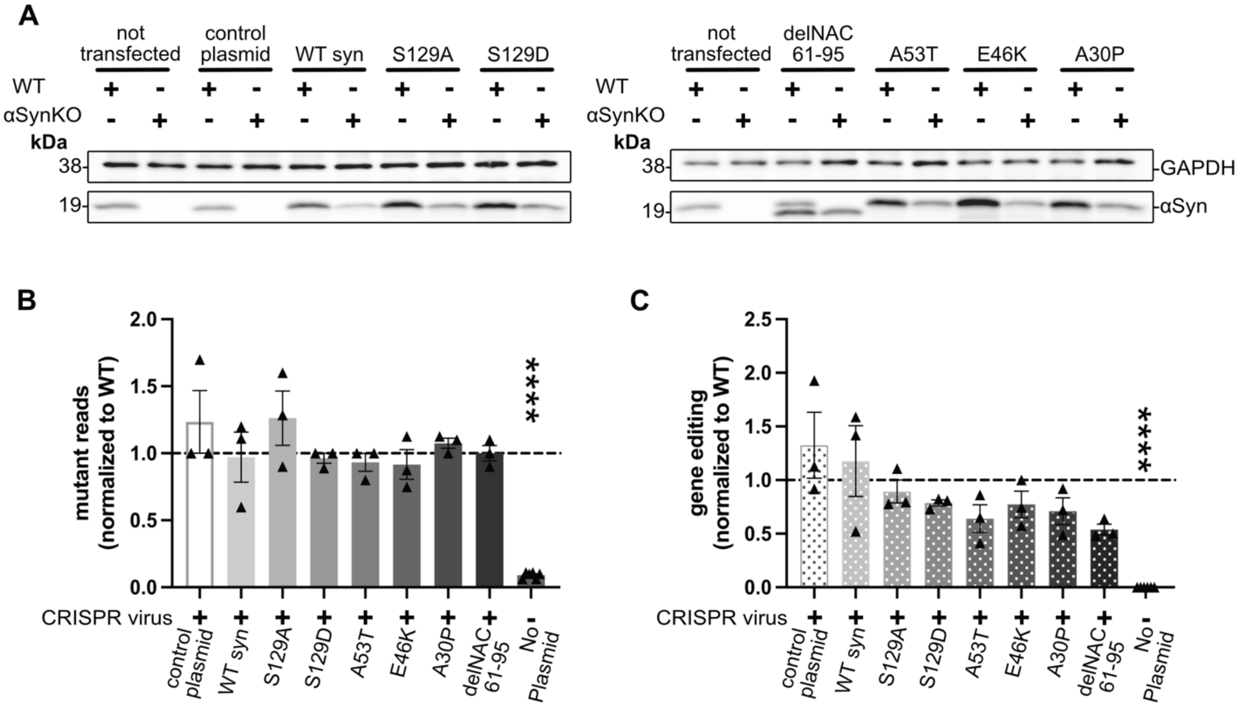


**Supplementary Figure 2. Transient transfection cannot be used to assay DSB repair in αSyn KO HAP1 cells.**

A) Western Blot post transfection confirming over-expression in WT HAP1 cells and full rescue in αSyn KO cells. Full rescue observed with WT syn, S129A syn, S129D syn, A53T syn, E46K syn, A30P syn, and delNAC 61-95 syn. B) Quantification of NGS of 288bp repair junction from αSyn KO cells with each synuclein construct condition normalized WT control plasmid. ANOVA summary p<0.0001. Post-hoc multiple comparisons show no significant differences. αSyn KO control plasmid (1.233 ±0.233) compared to WT control plasmid (p=0.5396). αSyn KO WT syn (0.9704±0.187)(p>0.9999). αSyn KO S129A (1.262 ±0.202)(p=0.4137). αSyn KO S129D (0.9630 ±0.037)(p>0.9999). αSyn KO A53T (0.9333 ±0.067)(p=0.9996). αSyn KO E46K (0.9167 ±0.110)(p-0.9977). αSyn KO A30P (1.075 ±0.038)(p=0.9990). αSyn KO delNAC 61-95 (1.000 ±0.058)(p>0.9999). αSyn KO Nontargeting virus (0.09120 ±0.009)(p<0.0001) N=3 biological replicates. One-way ANOVA. B) T7 Endonuclease I enzymatic assay quantification of percent gene editing of WT and αSyn KO cells transfected with WT and mutant forms of synuclein. % Gene Editing = 100 x (1 – (1-fraction cleaved)$^{1/2}$). ANOVA summary p<0.0001. Post-hoc multiple comparisons show no significant differences. αSyn KO control plasmid (1.325 ±0.308) compared to WT control plasmid (p=0.4784). αSyn KO WT syn (1.177 ±0.107)(p=0.9451). αSyn KO S129A (0.8936 ±0.107)(p=0.9982). αSyn KO S129D (0.7871 ±0.029)(p=0.8634). αSyn KO A53T (0.6414 ±0.129)(p=0.3716). αSyn KO E46K (0.7729 ±0.124)(p=0.8215). αSyn KO A30P (0.7102 ±0.125)(p=0.6015). αSyn KO delNAC61-95 (0.5394 ±0.050)(p=0.1494). αSyn KO Nontargeting virus (0.0 ±0.0)(p<0.0001). N=3 biological replicates. One-way ANOVA.

**Supplemental Figure 3.**

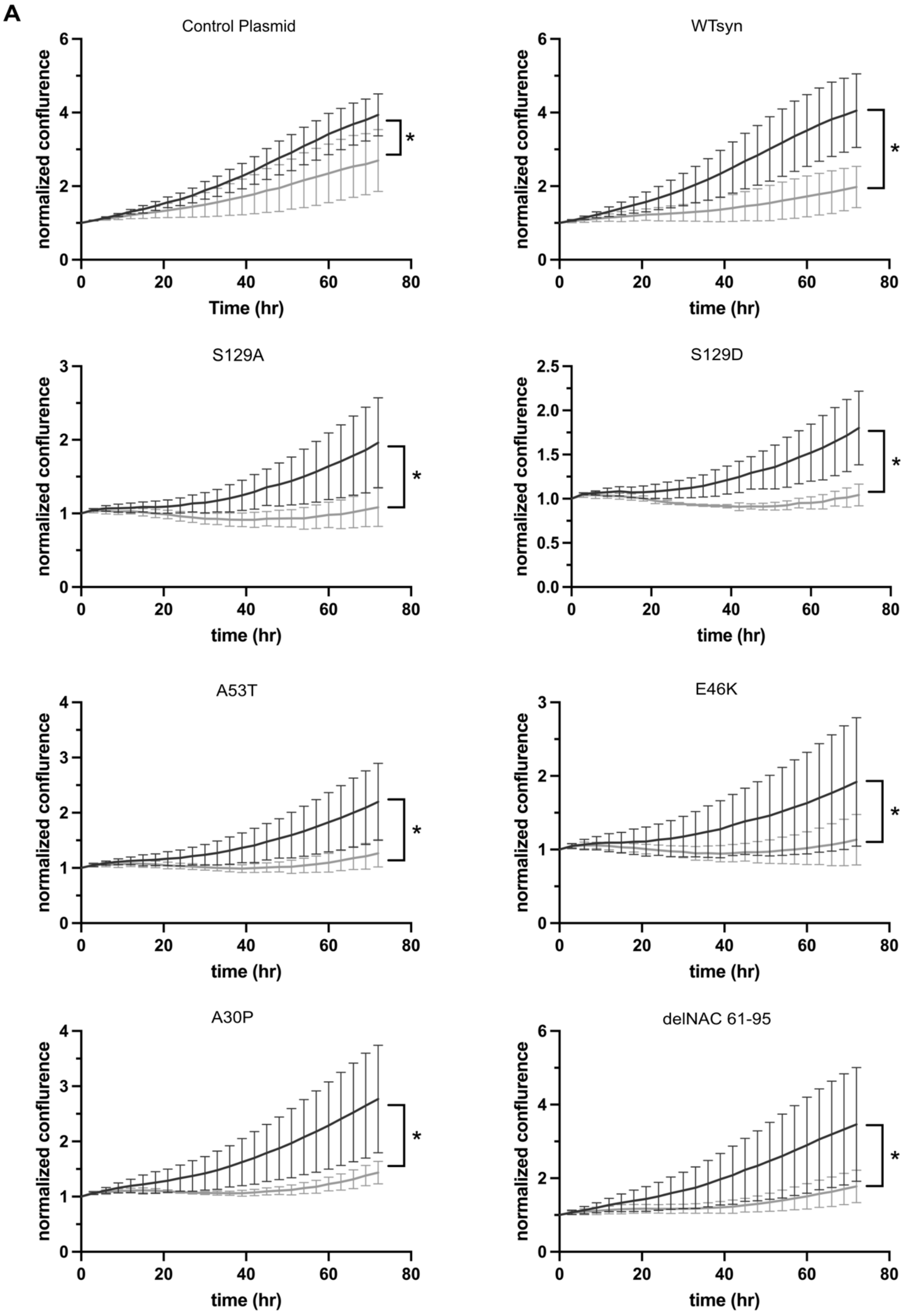

**Supplementary Figure 3. Re-expression of WT and mutant forms of αSyn does not affect proliferation of HAP1 cells.**

A) Proliferation assay of WT(black) and αSyn KO(gray) HAP1 cells transfected with WT and mutant forms of synuclein from Supplementary Figure 2. Significant differences between proliferative curves over 72 hours of cells transfected with control plasmids and mutants. WT control plasmid= 95% CI slope (0.03694-0.04849), KO control plasmid= 95% CI slope (0.01550- 0.03172). WT syn= 95% CI slope (0.03880-0.05068), KO WT syn= 95% CI slope (0.009200-0.01575). WT S129A= 95% CI slope (0.009390-0.01503), KO S129A= 95% CI slope (-0.001479-0.0009383). WT S129D= 95% CI slope (0.007822-0.01201), KO S129D= 95% CI slope (-0.001430– -0.0001155). WT A53T= 95% CI slope (0.01200-0.01867), KO A53T= 95% CI slope (0.0004279-0.002834). WT E46K= 95% CI slope (0.007699-0.01610), KO E46K= 95% CI slope (-0.001203-0.001774). WT A30P= 95% CI slope (0.01876-0.02842), KO A30P= 95% CI slope (0.002622-0.004776). WT delNAC61-95= 95% CI slope (0.02649-0.04193), KO delNAC61-95= 95% CI slope (0.006612-0.01099). N=3 biological replicates (6 technical replicates per biological replicate). Simple linear regression.

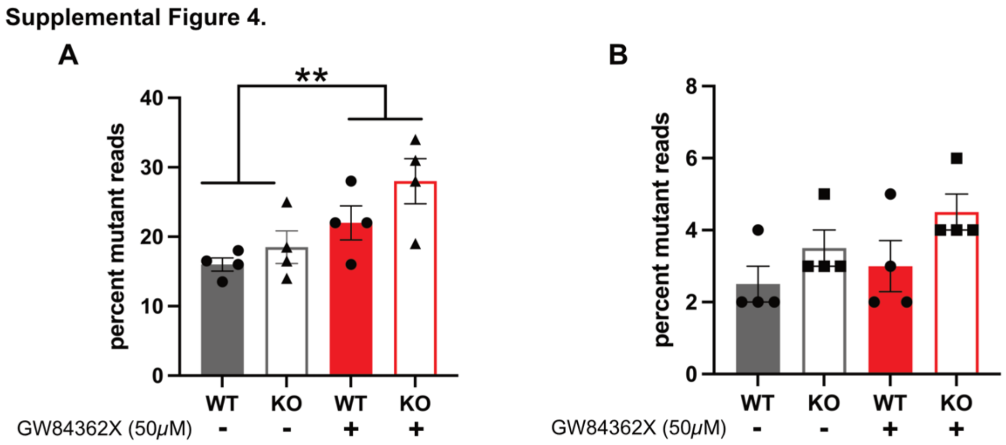


**Supplementary Figure 4: PLK1/3 inhibition increases indel frequency in HAP1 cells, but not in mouse cortical neurons.**

A) NGS of 288bp repair junction from WT and αSyn KO HAP1 cells transduced with CRISPR/Cas9 DSB inducing lentivirus and treated with 0.01% DMSO or PLK1/3 Inhibitor GW84362X 50μM for 72 hours. GW84362X treated cells show significantly increased mutant reads compared to DMSO treated cells. Row (Drug) Factor p=0.0071. Column (Cell Line) Factor p=0.1011. N=4 biological replicates (1-2 technical replicates per biological replicate). Two-way ANOVA. No post-hoc multiple comparisons. B) NGS of 272bp repair junction from WT and αSyn KO E18 mouse cortical neurons transduced with CRISPR/CAs9 DSB inducing lentivirus and treated with 0.01% DMSO or PLK1/3 Inhibitor GW84362X 50μM for 72 hours. Row (Drug) Factor p=0.2046. Column (Cell Line) Factor p=0.0451. N=4 biological replicates (1-2 technical replicates per biological replicate). Two-way ANOVA. No post-hoc multiple comparisons.
